# Intrinsic base substitution patterns in diverse species reveal links to cancer and metabolism

**DOI:** 10.1101/758540

**Authors:** Suzana P. Gelova, Kassidy N. Doherty, Salma Alasmar, Kin Chan

## Abstract

Analyses of large-scale sequencing data reveal that mutagenic processes create distinctive patterns of base substitutions, called mutational signatures. Here, we analyzed substitution patterns from seven model species and single nucleotide polymorphisms (SNPs) in 42 species, totalling >1.9 billion variants. We found that the base substitution patterns for many species most closely match mutational signature 5 in cancers. Signature 5 is also ubiquitous in cancers and normal human cells, suggesting similar patterns of mutation across species are likely due to conserved biochemistry. Finally, we present evidence from yeast that sugar metabolism is directly linked to this form of DNA damage. We propose that conserved metabolic processes in cells are coupled to continuous generation of mutations, which are acted upon by genetic selection to drive the evolution of species, and cancers.

**One Sentence Summary:** Energy metabolism produces DNA damage leading to similar patterns of base substitutions in many species and in human cancers.

## Main Text

Mutations in genomic DNA are the starting material for evolution by natural selection. Mutagenesis is often thought of as a stochastic process, wherein the temporal and positional occurrence of each mutational event is not predictable (*1*). And yet, sets of mutations induced by a given DNA damaging agent or process do create reproducible, characteristic patterns, called mutational signatures (*2*). Mutational signatures derived from cancer genomics data have been published, providing a composite picture of the recurrent patterns of base substitutions found in cancers, and in some cases, normal tissues as well (*3*). These observations in humans led us to investigate whether similar patterns of base substitutions are found when comparing genetic variation within and between other species.

We analyzed mutational data (784,102,480 single base substitutions) from seven model species, comprising: 1,011 varieties of budding yeast (*Saccharomyces cerevisiae*) (*4*); 161 varieties of fission yeast (*Schizosaccharomyces pombe*) (*5*); 152 isolates of roundworm (*Caenorhabditis elegans*) (*6*); 85 isolates of fruit fly (*Drosophila melanogaster*) (*7*); 36 strains of mouse (*Mus musculus*) (*8*); 42 strains of rat (*Rattus norvegicus*) (*9*); and 49 canines (*Canis familiaris*) (*10*). When the data were plotted using the convention for reporting mutational signatures (i.e., six substitution types, each subdivided into 16 contexts with one flanking base on either side, 5ʹ and 3ʹ) (*11*), it became clear that the majority of mutations in all seven species were either C→T or T→C transitions (see Figure 1A). To assess the similarity of these base substitution patterns to known mutational signatures, we computed cosine similarity values vs. a set of published COSMIC (<u>C</u>atalog <u>o</u>f <u>S</u>omatic <u>M</u>utations in <u>C</u>ancer) signatures (see Figure 1B) (*2*). For context, a cosine similarity value of 1 is obtained when comparing identical signatures. The substitution patterns for all seven model species were each most similar to COSMIC signature 5 (see Figure 1C, referred to hereafter as COSMIC 5), which is one of two ‘clock-like’ mutational signatures found in human, as the numbers of COSMIC 5 mutations essentially always increase monotonically with age (*12*). COSMIC 5 also accounts for ∼75% of de novo mutations in healthy human children (*13*). These results show that, just as COSMIC 5 is found ubiquitously in human, intrinsic base substitution patterns similar to COSMIC 5 are found in multiple model species.

**Figure 1.**
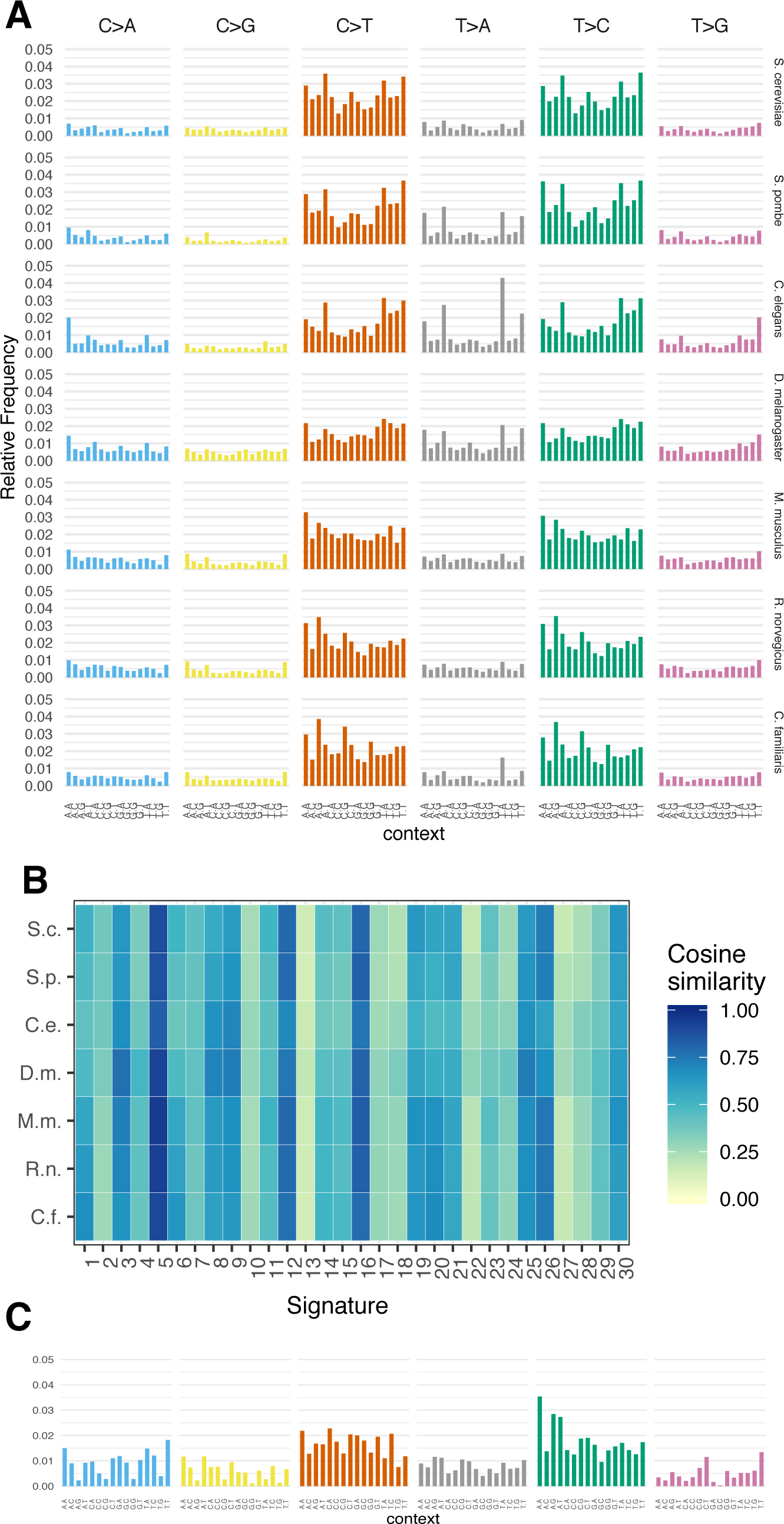
Intrinsic base substitution patterns of seven model species resemble one another and COSMIC 5. (A) 96-bar graphs showing the intrinsic base substitution patterns of seven model species, derived by analysis of data from (*4–10*). The patterns tended to show mostly C→T or T→C substitutions. Interestingly, both invertebrate species and fission yeast also had relatively prominent T→A peaks at ATT and TTA motifs. (B) Heatmap showing cosine similarity values to 30 COSMIC signatures. For all seven species, the highest cosine similarity values suggested closest resemblance to COSMIC 5. (C) COSMIC signature 5.

Given that COSMIC 5-like base substitution patterns are intrinsic to these model species, we then investigated if similar patterns are observed among SNPs in a larger, more diverse set of species. We downloaded sets of available VCF (variant call format) files from the NCBI SNP database for 44 non-human species (*14*). Two data sets were obviously invalid, as they completely lacked two substitution types (C→G and T→A, likely due to data processing error(s)), while another data set was dropped from analysis because of probable sequencing artefacts (see Figure S1). Among the remaining 41 species (yielding 603,722,210 SNPs, see Figure S2), the large majority showed the highest cosine similarity to COSMIC 5 (see Figure 2). We also analyzed 554,168,376 human SNPs from NCBI and found that they produced a pattern very similar to COSMIC 5 as well (cosine similarity = 0.957, see Figures 3A and S3). Taken altogether, observations from prior studies in human and our results here strongly suggest that conserved biochemical processes in many species are responsible for the set of COSMIC 5-like intrinsic base substitution patterns.

**Figure 2.**
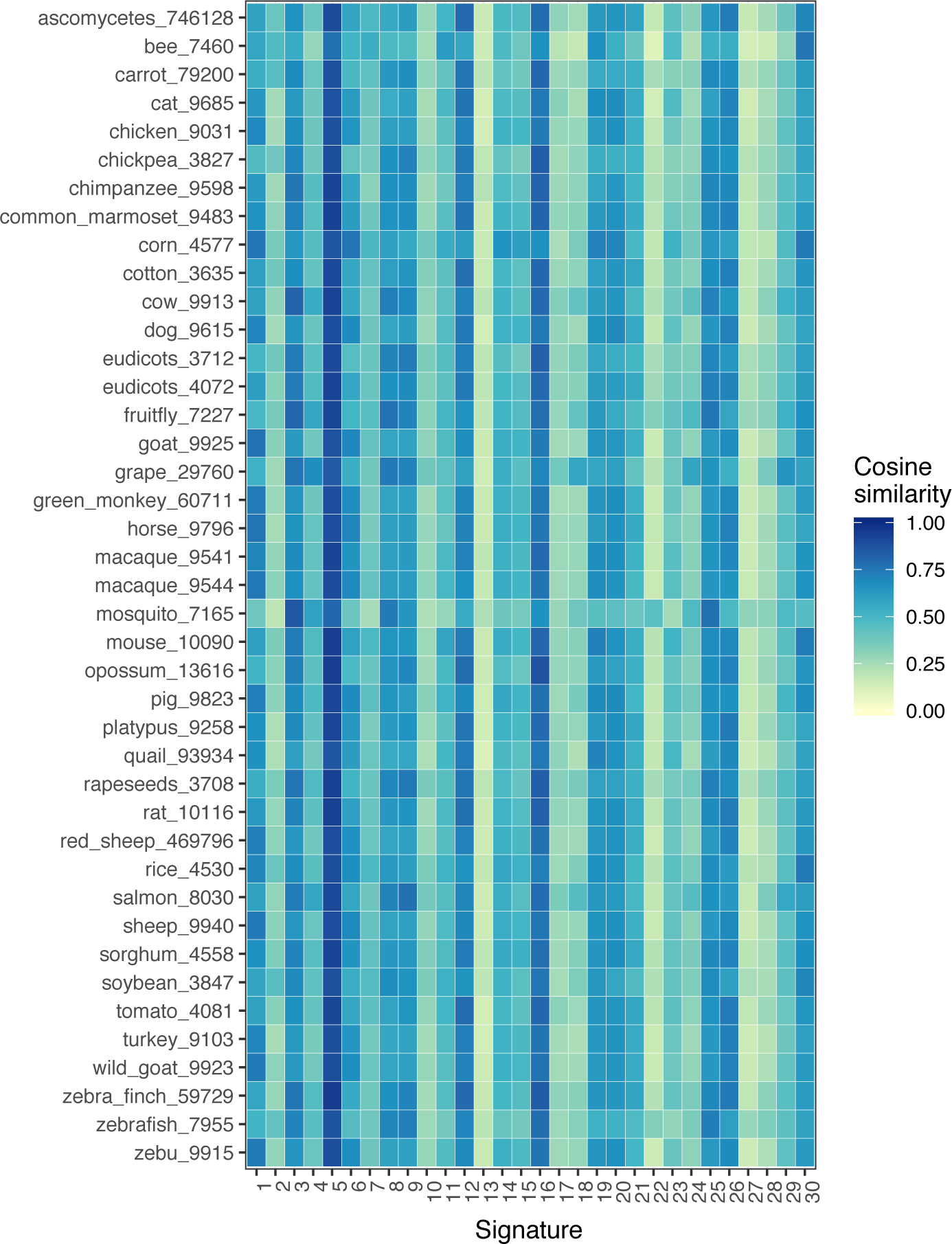
Heatmap depicting cosine similarity values comparing sets of SNPs from 41 non-human species vs. the 30 COSMIC signatures. Most species had the highest cosine similarity values compared to COSMIC signature 5. See Figure S2 for the 96-bar substitution patterns of these species.

**Figure 3.**
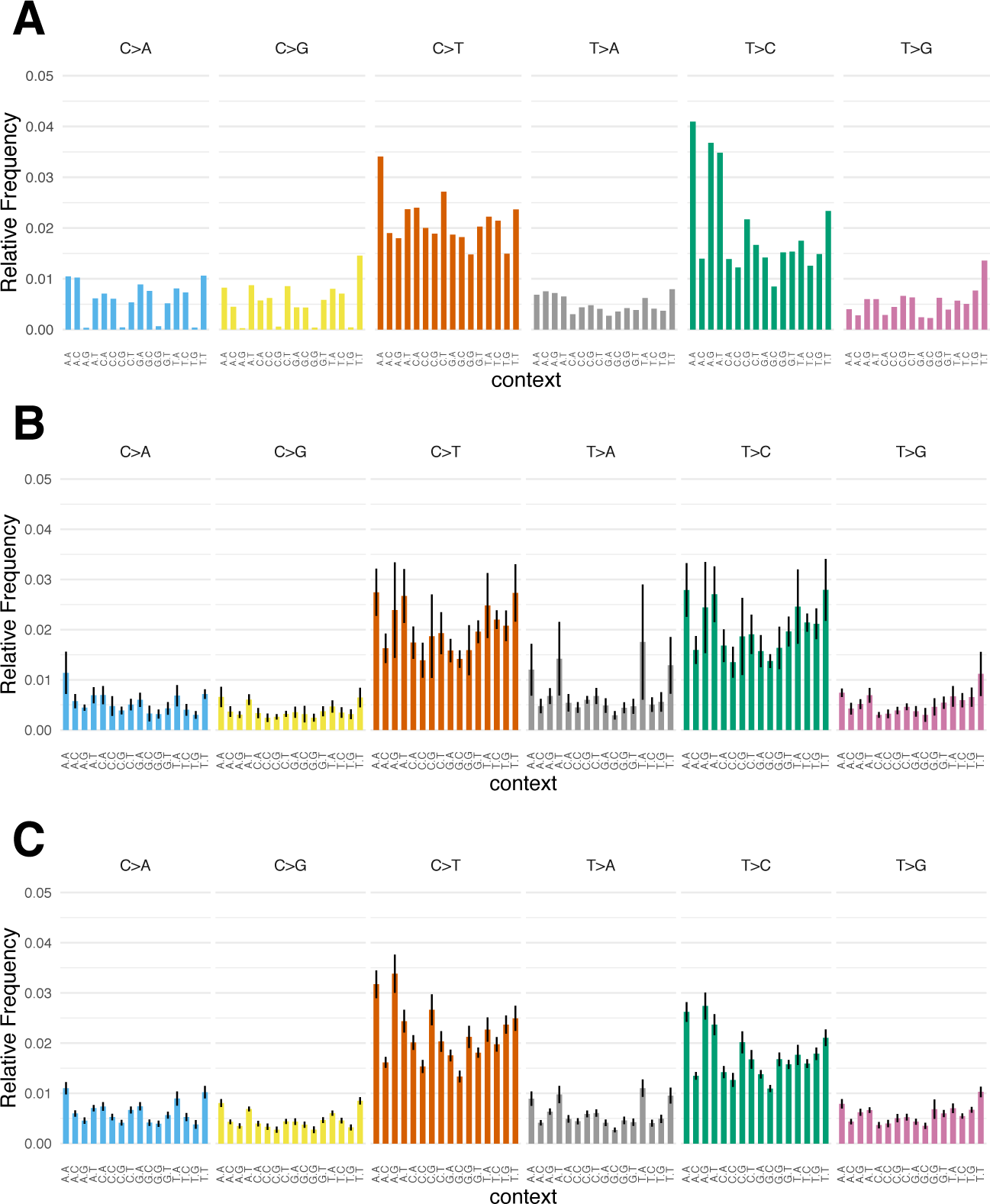
Base substitution patterns for NCBI human SNPs, model organisms, and NCBI SNPs from 41 non-human species. (A) The pattern obtained by plotting the entire collection of human SNPs from NCBI. (B) 96-bar graph showing mean and 95% confidence interval for relative frequencies of each substitution type among the seven model species. (C) 96-bar graph showing mean and 95% confidence interval for relative frequencies of each SNP type among the 41 non-human species from NCBI.

To quantify and visualize the extent of similarity among base substitution patterns, we generated profiles showing means and 95% confidence intervals of relative frequency for each mutation type among the seven model species (see Figure 3B). The 95% confidence intervals for most substitution types were relatively small, i.e., indicating low variance (see Figure S4A). The most highly variable trinucleotides with respect to differences in frequencies of mutations were TTA and ATT for T→A transversions (see also Figure 1A). Despite these instances of (relatively) higher variability, the mean values do provide a consensus COSMIC 5-like pattern among the seven model species. The cosine similarity between this consensus pattern and COSMIC 5 itself is 0.928 (see Figure S5).

Likewise, we applied this analysis to the SNP data from the NCBI database (see Figure 3C). The 95% confidence intervals for the means were more compact, reflecting the larger number of species examined and yielding a lower variance. Again, the substitution types with higher mean relative frequencies showed more variability (see Figure S4B). The consensus pattern from SNP data very closely matched the corresponding pattern from mutations among the seven model organisms, with a cosine similarity of 0.981 (see Figure S6). All four NCG motifs exhibit more common C→T transitions in the SNP data, suggestive of possible contributions from deamination of 5- methylcytosine at those motifs in a subset of mammals (e.g., goat, horse, macaque, sheep, etc.), similar to COSMIC signature 1 (*3*). Also similar to the model organism data, the consensus SNP pattern has a cosine similarity of 0.931 compared to COSMIC 5 (see Figure S7). Taken together, we propose that for many species, their respective sets of intrinsic base substitutions form a range of related patterns, and COSMIC 5 is simply the corresponding human counterpart in this continuum. The close relatedness of COSMIC 5 and the human SNPs pattern (see Figure S3) is consistent with this model.

In principle, many species can be used as models to study the molecular mechanisms underlying these COSMIC 5-like mutational patterns, as these mechanisms are likely to be well conserved. We previously used a budding yeast system which features controlled generation of genomic single-stranded DNA (ssDNA) to elucidate the relative contributions of APOBEC3A and APOBEC3B cytidine deaminases to cancer mutagenesis (*15*). This system relies on the *cdc13-1* temperature sensitive mutation (*16*), such that shifting to 37°C causes telomere uncapping, extensive DNA end resection, and cell cycle arrest in G_2_ via the DNA damage checkpoint (*17*). Since exposed ssDNA is much more prone to damage than double-stranded DNA, this system can acquire mutations for analysis much more readily than more conventional approaches. When we sequenced the genomes of cells that were allowed to accumulate mutations, we obtained a pattern of base substitutions that resembled the consensus base substitution pattern in the seven model species, the consensus pattern from SNPs of 41 species, and COSMIC 5 (cosine similarities = 0.941, 0.921, and 0.906, respectively, see Figures 4A and Figure S8-S10). This corroborates the generality of COSMIC 5-like intrinsic base substitution patterns.

**Figure 4.**
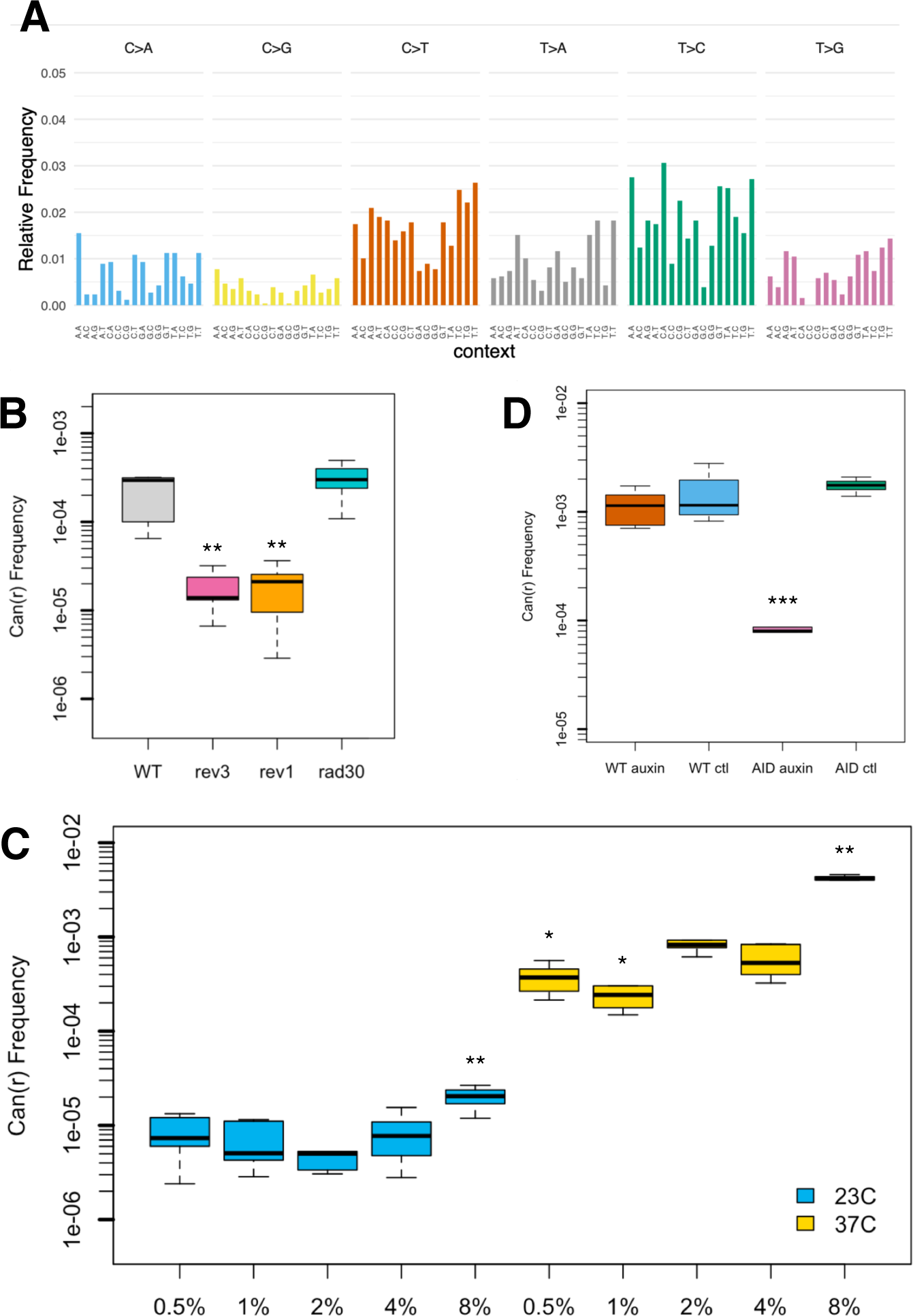
Corroborating genetic and genomic data from yeast model system. (A) Base substitution pattern resulting from damage to exposed single-stranded DNA in yeast metabolizing glucose. This is most closely related to the model species consensus substitution pattern in Figure 3B, with cosine similarity of 0.941. (B) *CAN1* reporter gene inactivation frequency was dependent on *REV3* (encoding the catalytic subunit of polymerase zeta) and *REV1*, i.e., on error-prone translesion DNA synthesis. Deletion of *RAD30* (encoding polymerase eta) had no effect. Cells were temperature shifted to 37°C for 6 hours. (C) Increasing glucose concentration to 8% resulted in a significant increase in mutagenesis within single-stranded DNA, when compared to baseline conditions in 2% glucose. Cells in 0.5% and 1% glucose both experienced significantly less mutagenesis than controls in 2%. Cells were shifted to 37°C for 24 hours. (D) Mutagenesis was dependent on completion of glycolysis. Inducible degradation of Cdc19 (pyruvate kinase) blocked the last step of the glycolytic pathway and caused over a ten-fold decrease in mutagenesis. Cells were shifted to 37°C for 24 hours in the presence or absence of 1 mM auxin. * denotes *P* < 0.05; ** is *P* < 0.01; and *** is *P* < 0.001 by t- test.

We then carried out additional experiments to investigate mechanisms responsible for the mutational pattern. Mutagenesis is dependent on the translesion DNA synthesis (TLS) polymerases zeta and Rev1 (*18*), i.e., on the error-prone bypass of damaged bases (*19*) (see Figure 4B). In contrast, deletion of *RAD30* (encoding polymerase eta) had no effect, since Rad30 specializes in error-free bypass of ultraviolet light-induced cyclobutane dimer photoproducts (*20*) and apparently plays no role in mutagenesis opposite DNA damage that gives rise to the COSMIC 5-like pattern. Deletion of genes important for error-free bypass also had no significant effect (see Figure S11). Moreover, we found that energy metabolism is key to this form of mutagenesis. Cells maintained in glucose (a fermentative, glycolytic energy source (*21*)) acquired mutations most frequently, while cells in acetate (a carbon source that can be introduced directly into the tricarboxylic acid cycle, and must be utilized by aerobic respiration (*22*)) had the fewest acquired mutations (see Figure S12). Moreover, increasing the concentration of glucose from 2% (the amount normally used in yeast media) to 8% led to a five-fold increase in the frequency of inactivating mutations in the *CAN1* reporter gene (see Figure 4C).

Blocking the last step of glycolysis using auxin-induced degron alleles of *CDC19* (encoding pyruvate kinase) caused a large decrease in mutagenesis (see Figure 4D), further supporting the central role of energy metabolism to generate the DNA damage which produces a COSMIC 5-like base substitution pattern. Finally, the pattern of mutations acquired in 8% glucose is essentially the same pattern as in 2% glucose (cosine similarity = 0.970, see Figure S13); there is simply more DNA damage overall in 8% glucose.

Taken altogether, our results suggest the intrinsic base substitution patterns of many organisms are linked to sugar metabolism. This would provide a plausible explanation for why such similar patterns are found across many species, including human, since the main pathways of sugar metabolism are well conserved (*23*). This model also implies that processes of sugar metabolism will inevitably produce DNA damage, which leads to mutations that fit a similar pattern across many species. In turn, these mutations are starting material for evolution by natural selection. Since energy generation is a prerequisite for living, the continuous basal level of mutagenesis which results would ensure that natural selection will have an ever-renewing repertoire of genetic variants to act upon. We propose the actual DNA damage itself is likely very similar between species, but the resulting patterns are more or less divergent depending on the similarity of the respective sets of error-prone TLS polymerases when comparing between species (see Figures S14 and S15). Species with polymerases that are more closely related would have more similar patterns than those from more diverged species. Finally, a coupling between metabolism and COSMIC 5-like mutational patterns could help explain why the extent of that type of mutagenesis can be highly variable in cancers, when comparing between different patients (*12*) or temporally within the same patient (*24*): namely, such variability would be linked to differences in sugar metabolism between individuals or over the course of disease progression in particular patients.

## Acknowledgments

We thank all members of the Chan laboratory and numerous other colleagues for their critical feedback on this work. Funding was provided by a Tier 2 Canada Research Chair, an NSERC Discovery Grant, an Ontario Early Researcher Award, and uOttawa startup funding to K.C. Author contributions are as follows: All authors performed experiments and discussed the data; S.P.G. and K.C. analyzed data; K.C. conceived and supervised the study, and wrote the manuscript. The authors declare there are no competing interests. Genomics data are available online as described in the Data Acquisition section within Materials and Methods. Yeast strains are available on request.

## Materials and Methods

### Data Acquisition

VCF files were downloaded from the following sources: *Saccharomyces cerevisiae*, http://1002genomes.u-strasbg.fr/files/; *Schizosaccharomyces pombe*, ftp://ftp.pombase.org/pombe/external_datasets/DJeffares_Diversity/; *Caenorhabditis elegans*, https://www.elegansvariation.org/data/release/20160408; *Drosophila melanogaster*, https://trace.ncbi.nlm.nih.gov/Traces/sra/sra.cgi?study=SRP050151; *Mus musculus*, ftp://ftp-mouse.sanger.ac.uk/REL-1807-SNPs_Indels/; *Rattus norvegicus*, ftp://ftp.rgd.mcw.edu/pub/strain_specific_variants/Hermsen_et_al_40Genomes_Variants/ ; *Canis familiaris*, ftp://download.big.ac.cn/dogsd/dog10k/variations/58indiv.unifiedgenotyper.recalibrate__95.5_filtered.pass_snp.vcf.gz; NCBI non-human SNP data, ftp://ftp.ncbi.nih.gov/snp/organisms/archive/; and human SNP data, ftp://ftp.ncbi.nih.gov/snp/latest_release/VCF/GCF_000001405.38.gz. Illumina WGS reads from ssDNA system in either 2% or 8% glucose are here: https://www.ncbi.nlm.nih.gov/sra/PRJNA608117.

### Mutational Analyses of Downloaded VCFs

Where possible, we used the MutationalPatterns R package (version 1.10.0) to extract mutational and trinucleotide contexts from VCF files, to plot mutational signatures, to calculate cosine similarities, and to plot heatmaps (*11*). We constructed BSgenome (*25*) reference genome packages in order to use MutationalPatterns to process VCF files, if none of the stock reference genomes were appropriate. In cases where an organism’s reference genome proved to be incompatible with BSgenome 1.52.0 (or more precisely, its dependency GenomeInfoDB 1.20.0 (*26*)), we used a script that extracted mutational and trinucleotide context data. This script used a combination of Unix shell functions, bcftools 1.9 (*27*), VCFtools 0.1.15 (*28*), base R (*29, 30*), and the MafTools 2.0.16 R package (*31*). Summed matrices of mutational counts were loaded into MutationalPatterns for graphics rendering, in order to produce a consistent style for figures. We noticed that sometimes, GenomeInfoDB was unable to resolve the appropriate reference genome. A workaround for this issue is to ensure that only one file, specifying the chromosome names of the species being analyzed, is present in the ∼/Library/R/3.6/library/GenomeInfoDb/extdata/dataFiles folder. When the irrelevant files for other species are moved (temporarily) to a different folder, GenomeInfoDB is not given the option to use an incorrect reference. Most analyses were run on a custom built PC operating Ubuntu version 18.04, with the remaining analyses run on a MacBook Pro operating macOS version 10.14.5 or 10.14.6. R version 3.6.0 was used on both computing platforms.

### Reagents and Consumables

Bacto peptone (product code 211677) and yeast extract (212750) were purchased from Becton, Dickinson and Co. (Franklin Lakes, New Jersey). Auxin (3-indoleacetic acid, I3750), canavanine (C9758), and adenine sulfate dihydrate (AD0028) were purchased from MilliporeSigma (St. Louis, Missouri). Agar (FB0010), carbon sources (i.e., glucose [GB0219], galactose [GB0215], raffinose [RJ392], and potassium acetate [PB0438]), hygromycin (BS725), PCR purification spin column kit (BS654), agarose (D0012), and Tris-Borate-EDTA (TBE) buffer (A0026) were purchased from BioBasic (Markham, Ontario). G418 sulfate (450-130) was purchased from Wisent (St-Bruno, Québec). Q5 PCR kit were purchased from New England Biolabs Canada (Whitby, Ontario). Plasmid pOsTIR1w/oGFP (for integrating the *Oryza sativa TIR1* gene into the *HO* locus in yeast) was purchased from Addgene (Watertown, Massachusetts). pMK152 plasmid (for degron tagging *CDC19*) was purchased from the National BioResource Project Yeast Genetic Resource Center (NBRP YGRC, Osaka, Japan). Petri dishes (82.1473.001) were purchased from Sarstedt (Montréal, Québec).

### Yeast Genetics

All yeast strains were constructed in the ySR127 background, a *MATα* haploid bearing the *cdc13-1* temperature sensitive allele. In addition, there is a cassette of three reporter genes (*CAN1*, *URA3*, and *ADE2*) near the de novo left telomere of chromosome V. Details about ySR127 were described previously (*32*).

Gene knockouts were constructed by one step gene replacement using antibiotic resistance marker cassettes (*33*). To construct the CDC19 degron strains, we first introduced the *O. sativa TIR1* gene cassette (OsTIR1) into the *HO* locus of ySR127, using the protocol described in (*34*). We then inserted the 3mAID degron tag (containing three tandem repeats of the core domain of the IAA17 protein from *Arabidopsis thaliana*), using the protocol described in (*35*). Genetic constructions were verified by diagnostic replica plating and PCR.

Mutagenesis experiments were initiated by inoculating single colonies each into 5 mL of rich media (2% Bacto peptone, 1% Bacto yeast extract, specified percentage of a carbon source, supplemented with 0.001% adenine sulfate) in round bottom glass tubes. Cells were grown at permissive temperature (23°C) for three days. Then, cultures were diluted ten-fold into fresh media and shifted to restrictive temperature (37°C). Auxin was added to 1 mM final concentration for degron experiments. After temperature shift, cells were collected by light centrifugation, washed in water, and plated (using a turntable and cell spreader) onto synthetic complete media to assess survival and onto canavanine- containing media to select for mutants. Further details of this procedure are described in (*36*).

### Illumina Whole Genome Sequencing

To generate mutants for sequencing, four replicate ySR127 colonies were each suspended into a small volume of water (e.g., 50 μL). Half of each cell suspension was used to inoculate 5 mL of YPDA liquid media (2% Bacto peptone, 1% Bacto yeast extract, 2% glucose, supplemented with 0.001% adenine sulfate). The other half was used to inoculate 5 mL of YPDA-8 (2% Bacto peptone, 1% Bacto yeast extract, 8% glucose, supplemented with 0.001% adenine sulfate). Cultures were grown for three days at 23°C. 500 μL of each culture was then diluted with 4.5 mL of fresh media and shifted to 37°C. Cells from the remainder of each 23°C culture were collected by centrifugation and pellets were frozen at -20°C. Can^r^ Ade^-^ mutants were collected and reporter gene loss of function phenotypes were verified as described (*36*). Genomic DNA from mutants of interest were extracted and sequenced as described (*15*). Illumina whole genome sequencing was outsourced to Genome Québec (McGill University, Montréal). Eight parental cultures (actually four pairs of matched 2% and 8% glucose cultures) and 87 mutants derived from those parentals were sequenced. Bowtie2 version 2.3.5.1 (*37*), SAMtools 1.9 (*38*), and bcftools 1.9 (*27*) were used to map reads and call variants.

Variants with quality score < 30 and/or with sequencing coverage < 10 were filtered out. Variants found in parentals were filtered from the sets of variants identified in the corresponding derivative mutants. The resulting VCF files, containing mutant-specific variants, were passed to MutationalPatterns for further analysis and visualization.

## Supplementary Figures

**Fig. S1.**
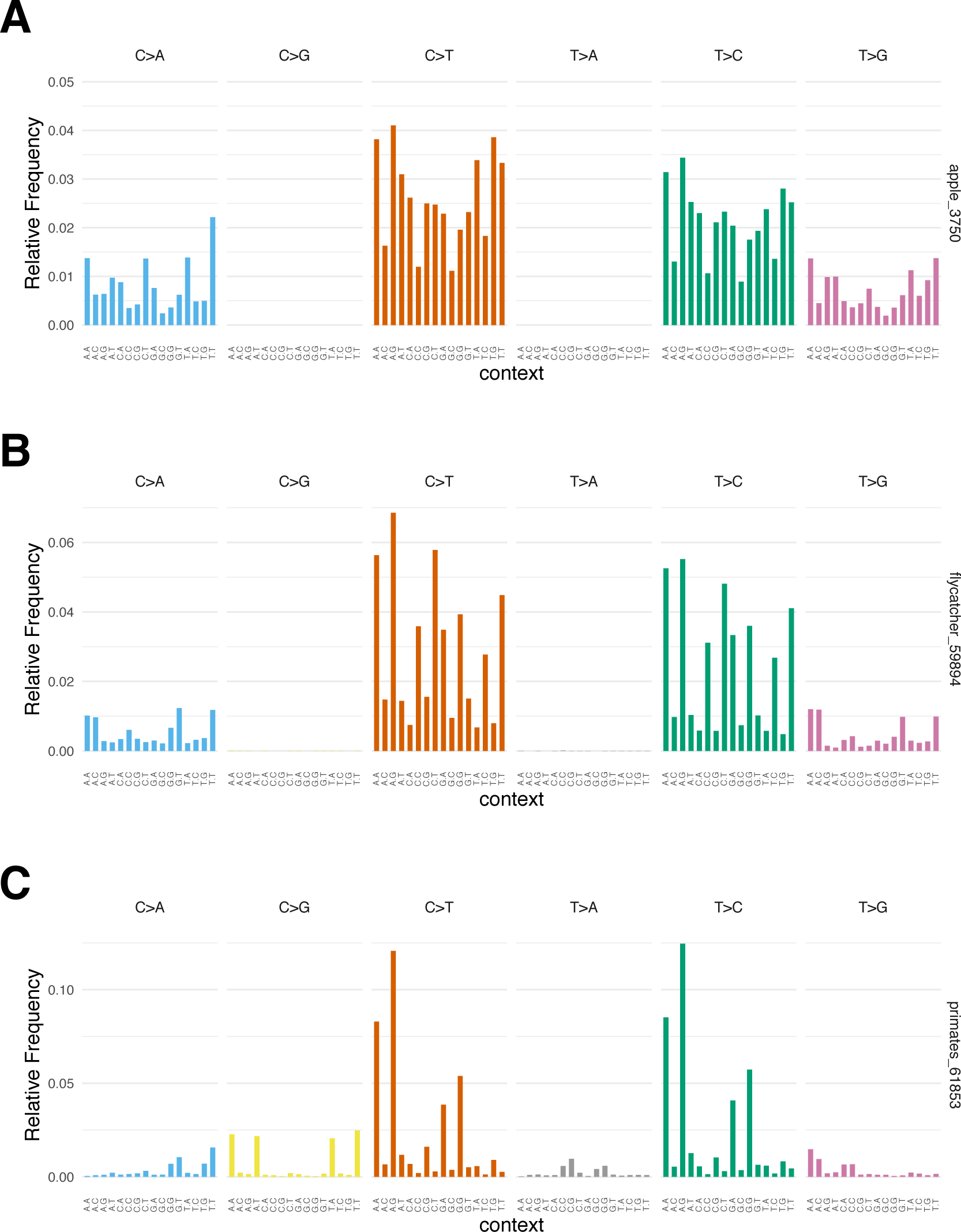
SNP patterns from NCBI database that were disqualified from further analysis. (A) Plotting of the SNP dataset for apple (*Malus domestica*) revealed an apparent data processing issue that deleted all C→G and T→A substitutions. (B) The SNP dataset for flycatcher (*Ficedula albicollis*) was also affected by this issue. (C) The SNP pattern for *Nomascus leucogenys* (the northern white-cheeked gibbon, but referred to simply as “primates” on NCBI) was also suspiciously unusual with very high peaks at the reciprocal motifs ACA and ATA, as well as ACG and ATG, for both C→T and T→C substitutions. As this was very much an outlier, the *Nomascus leucogenys* dataset was dropped for further consideration due to suspected sequencing artefact.

**Fig. S2.**
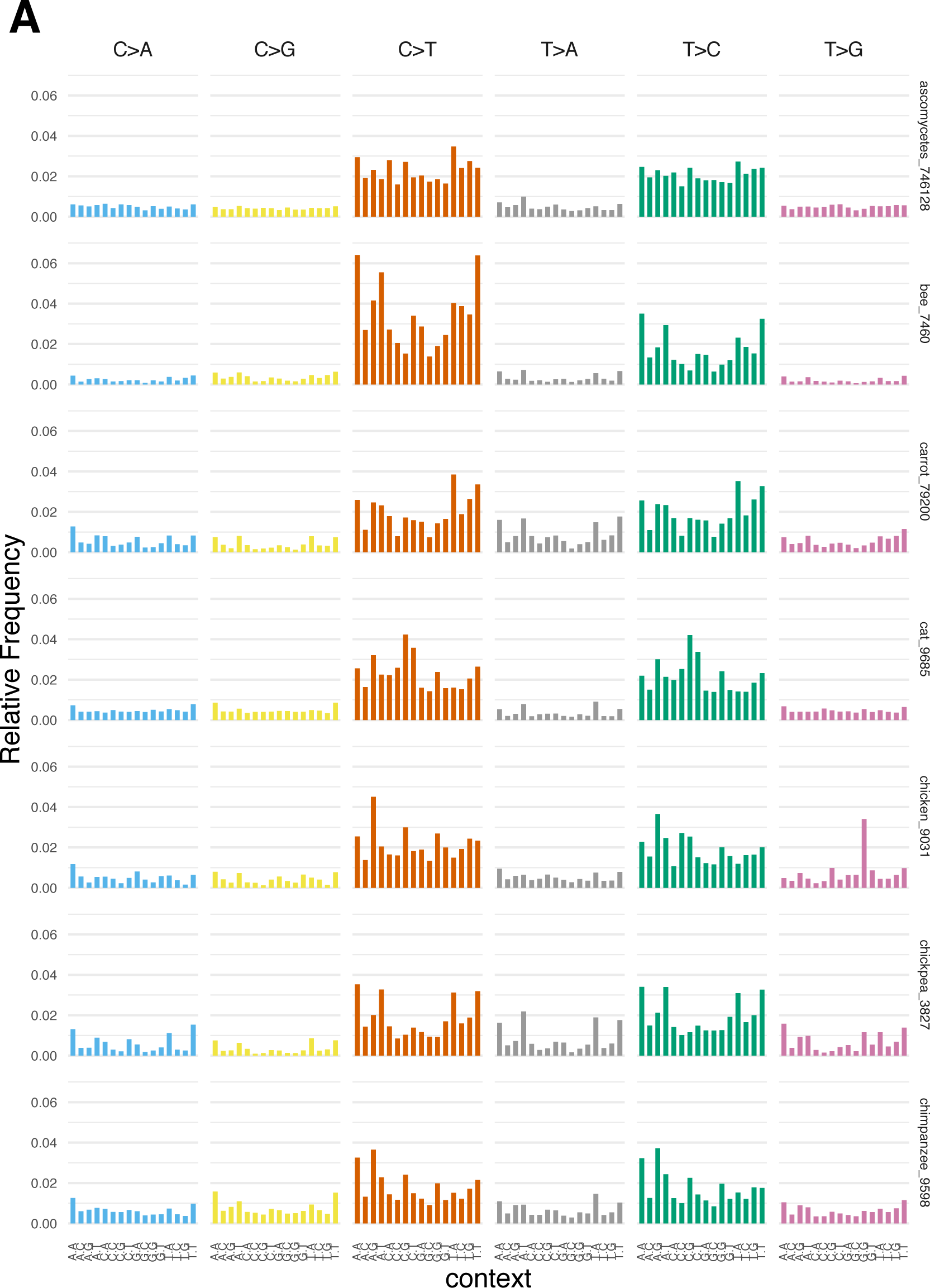
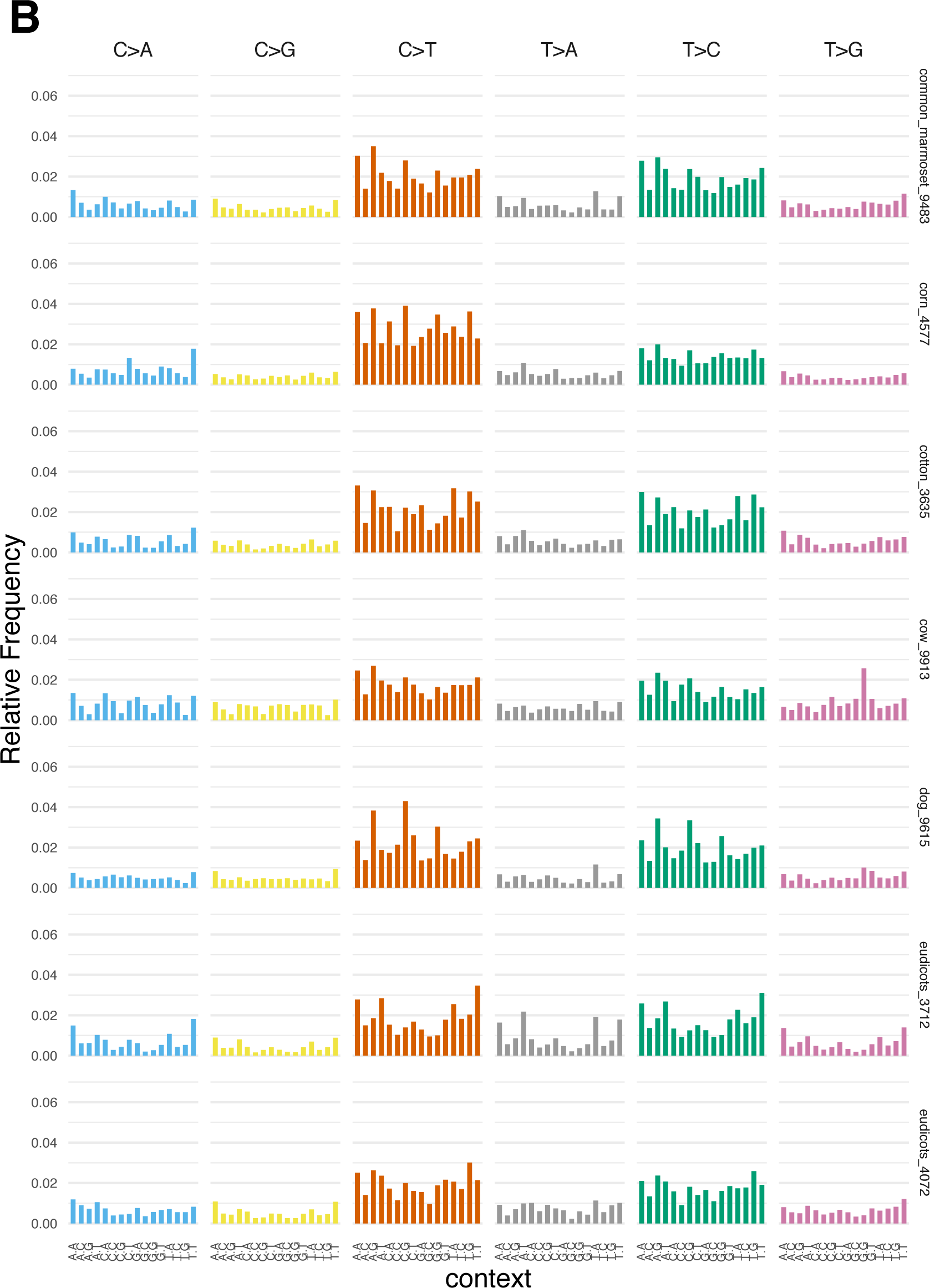
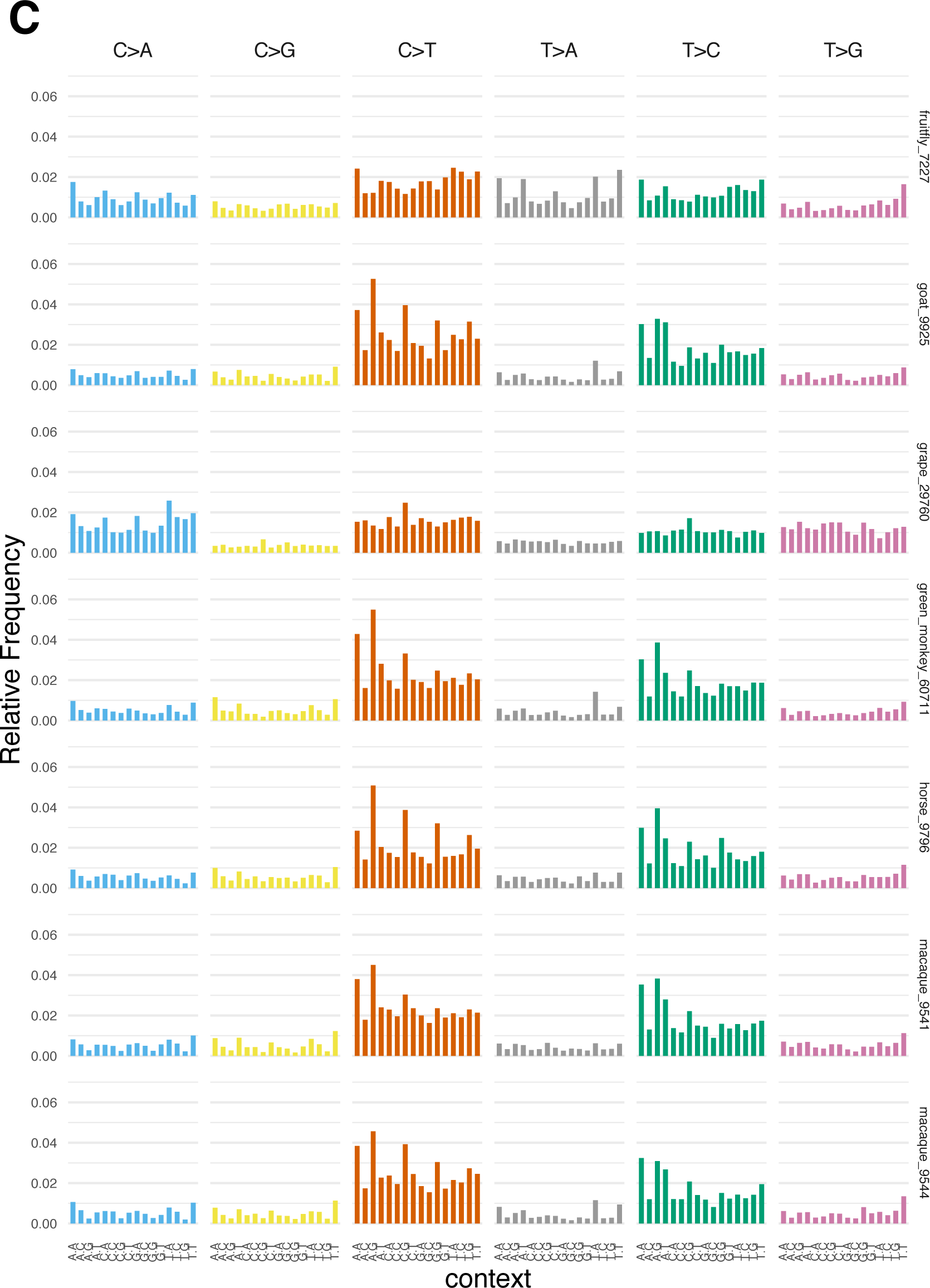
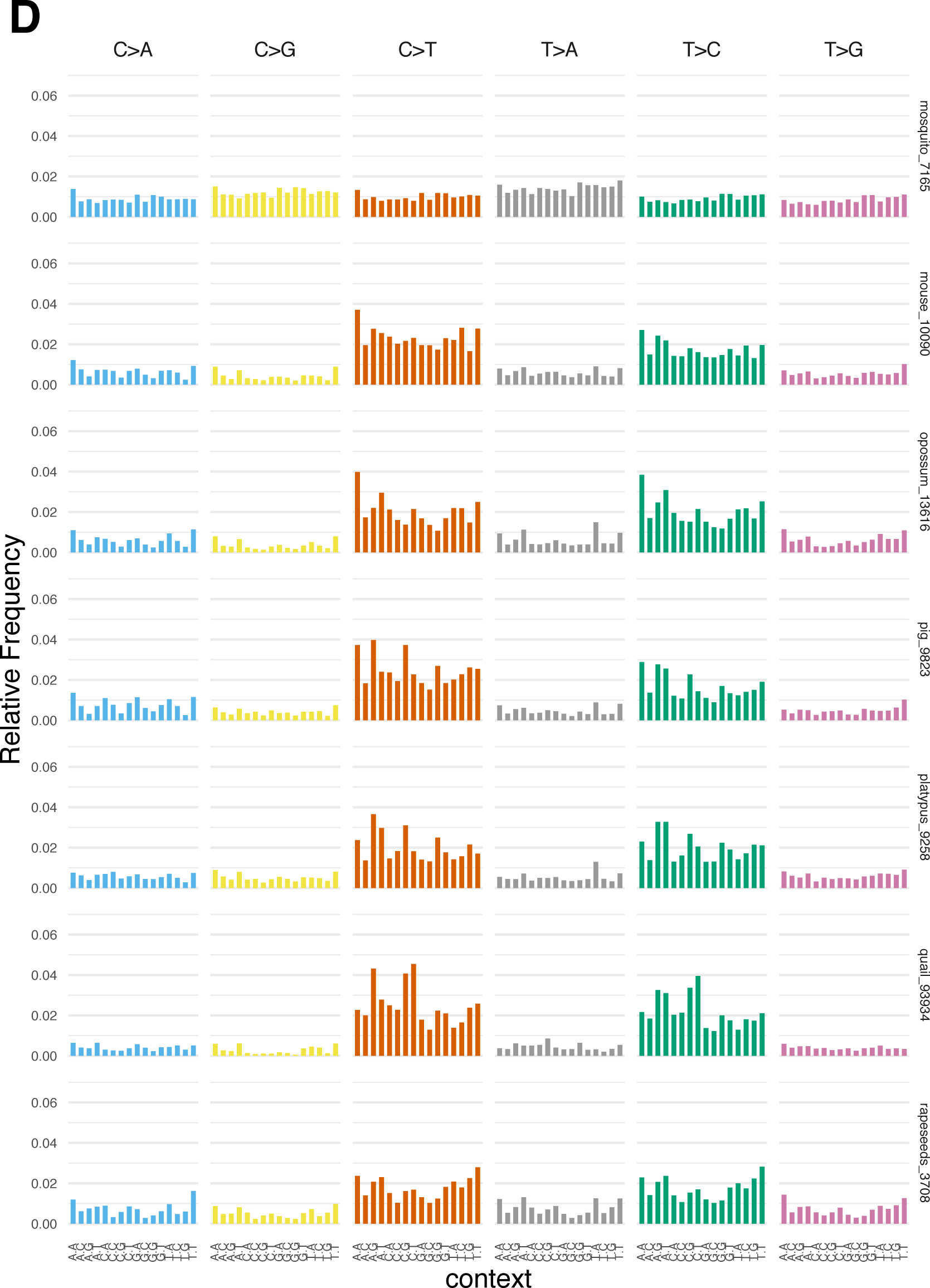
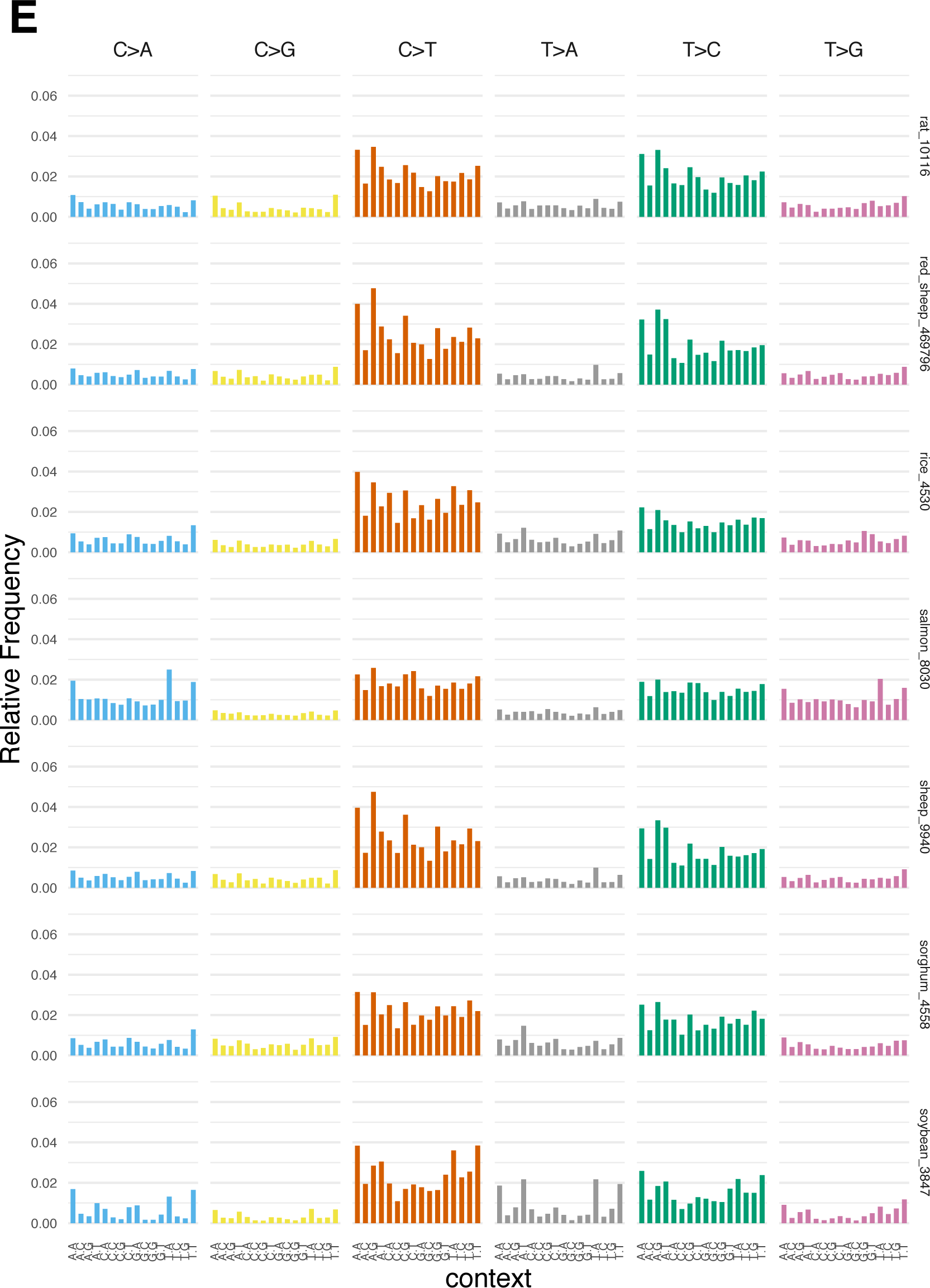
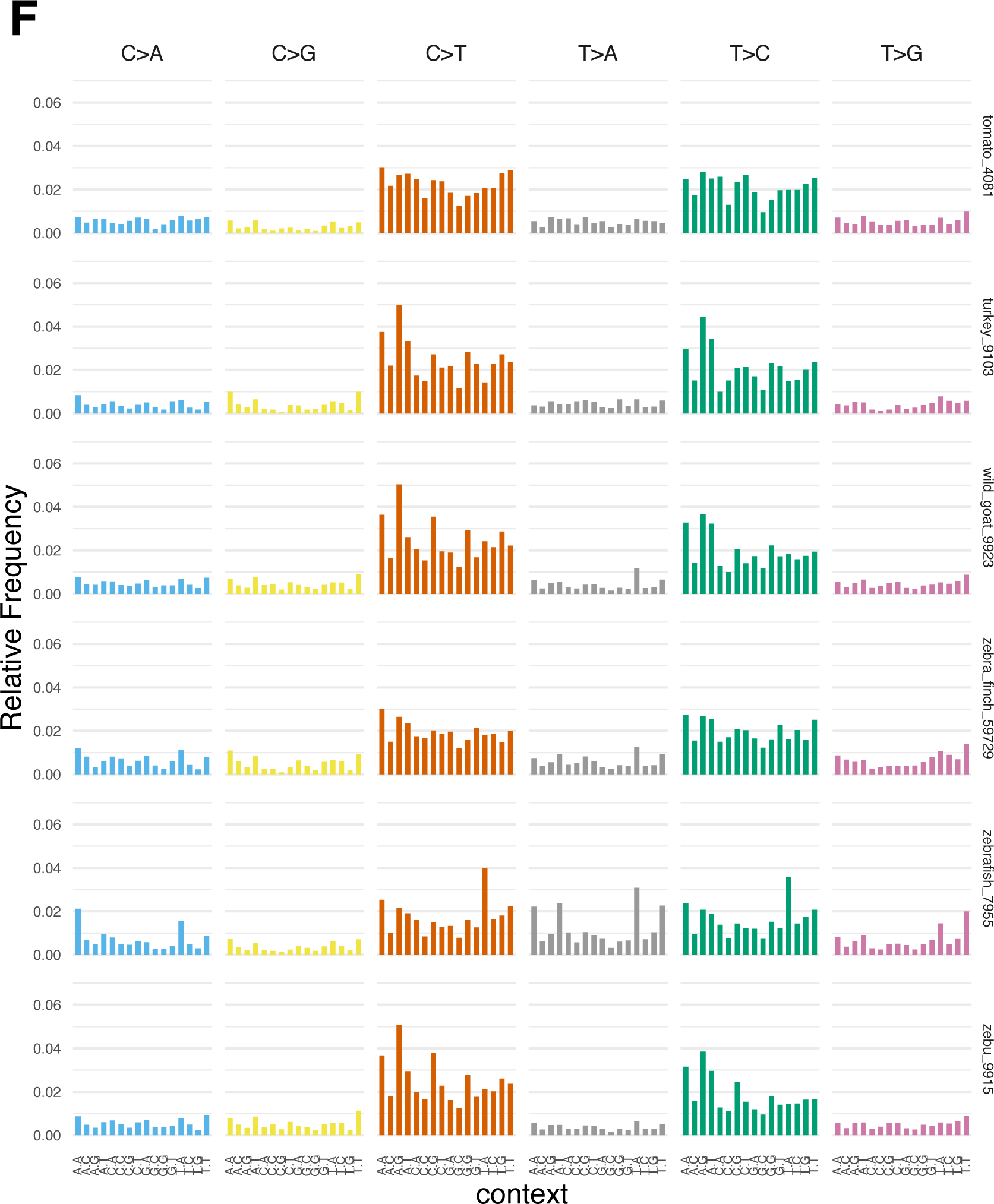
96-bar profiles showing the intrinsic base substitution patterns inferred from SNPs of 41 non-human species downloaded from NCBI. (A) Patterns for *Aspergillus fumigatus* (ascomycetes_746128), *Apis mellifera* (bee_7460), *Daucus carota* (carrot_79200), *Felis catus* (cat_9685), *Gallus gallus* (chicken_9031), *Cicer arietinum* (chickpea_3827), and *Pan troglodytes* (chimpanzee_9598). (B) Patterns for *Callithrix jacchus* (common_marmoset_9483), *Zea mays* (corn_4577), *Gossypium hirsutum* (cotton_3635), *Bos taurus* (cow_9913), *Canis familiaris* (dog_9615), and *Brassica oleracea* (eudicots_3712). (C) Patterns for *Drosophila melanogaster* (fruitfly_7227), *Capra hircus* (goat_9925), *Vitis vinifera* (grape_29760), *Chlorocebus sabaeus* (green_monkey_60711), *Equus caballus* (horse_9796), *Macaca fascicularis* (macaque_9541), and *Macaca mulatta* (macaque_9544). (D) Patterns for *Anopheles gambiae* (mosquito_7165), *Mus musculus* (mouse_10090), *Monodelphis domestica* (opossum_13616), *Sus scrofa* (pig_9823), *Ornithorhynchus anatinus* (platypus_9258), *Coturnis japonica* (quail_93934) and *Brassica napus* (rapeseeds_3708). (E) Patterns for *Rattus norvegicus* (rat_10116), *Ovis orientalis* (red_sheep_469796), *Oryza sativa* (rice_4530), *Salmo salar* (salmon_8030), *Ovis aries* (sheep_9940), *Sorghum bicolor* (sorghum_4558), and *Glycine max* (soybean_3847). (F) Patterns for *Solanum lycopersicum* (tomato_4081), *Meleagris gallopavo* (turkey_9103), *Capra hircus* (wild_goat_9923), *Taeniopygia guttata* (zebra_finch_59729), *Danio rerio* (zebrafish_7955), and *Bos indicus* (zebu_9915). Most patterns showed, to varying extents, a preponderance of C→T and T→C substitutions where these substitution types were similar in overall proportion, forming a range of COSMIC 5-like patterns. Notable exceptions to this overarching meta-pattern were mosquito, bee, grape, and corn.

**Fig. S3.**
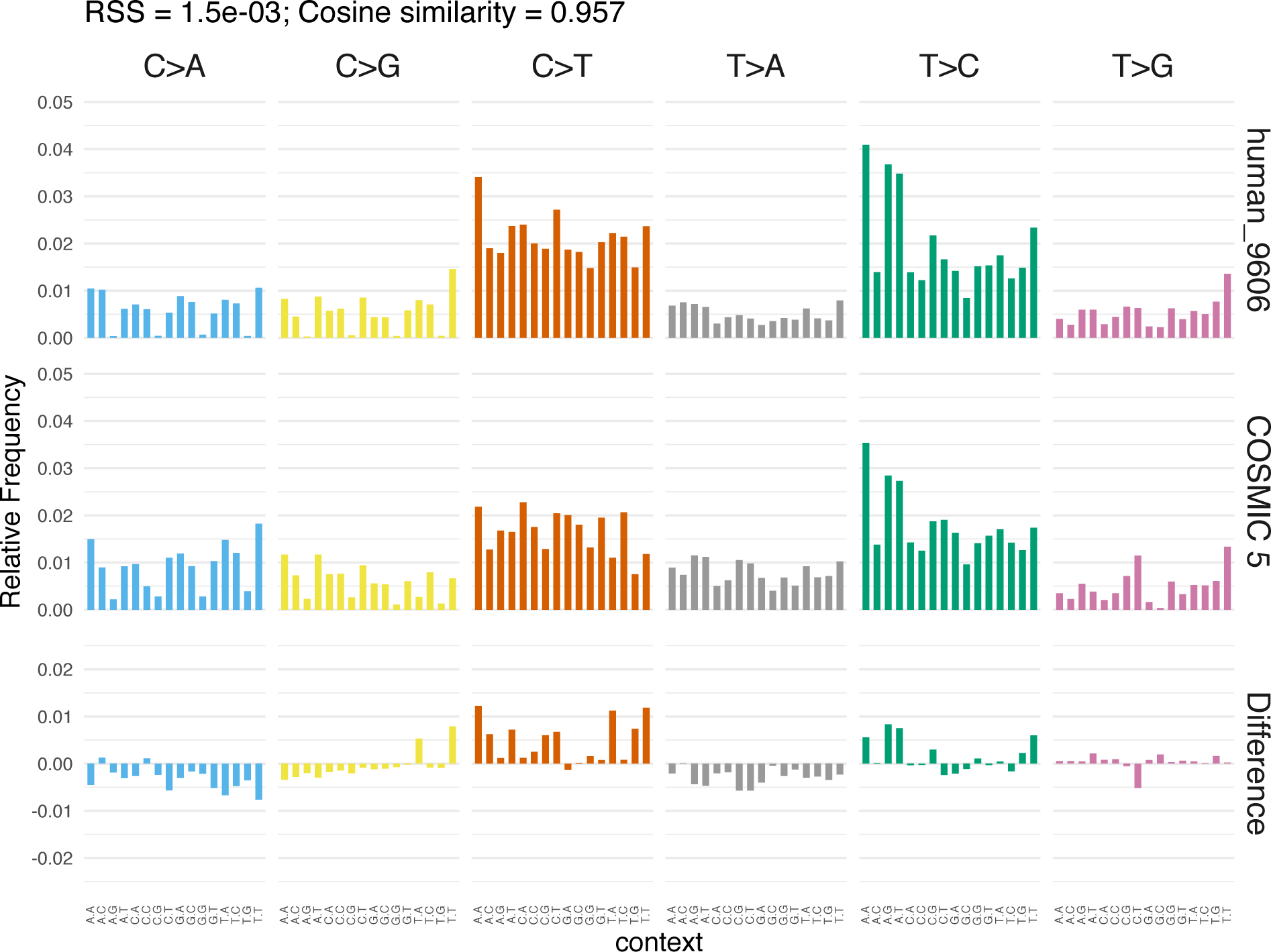
Comparison of the substitution profile from 554,168,376 human SNPs vs. COSMIC signature 5. The high cosine similarity value of 0.957 suggested that the mutagenic processes happening in cancers and over evolutionary time in humans are most likely the same, but the rate of mutation acquisition is significantly higher in cancers.

**Fig. S4.**
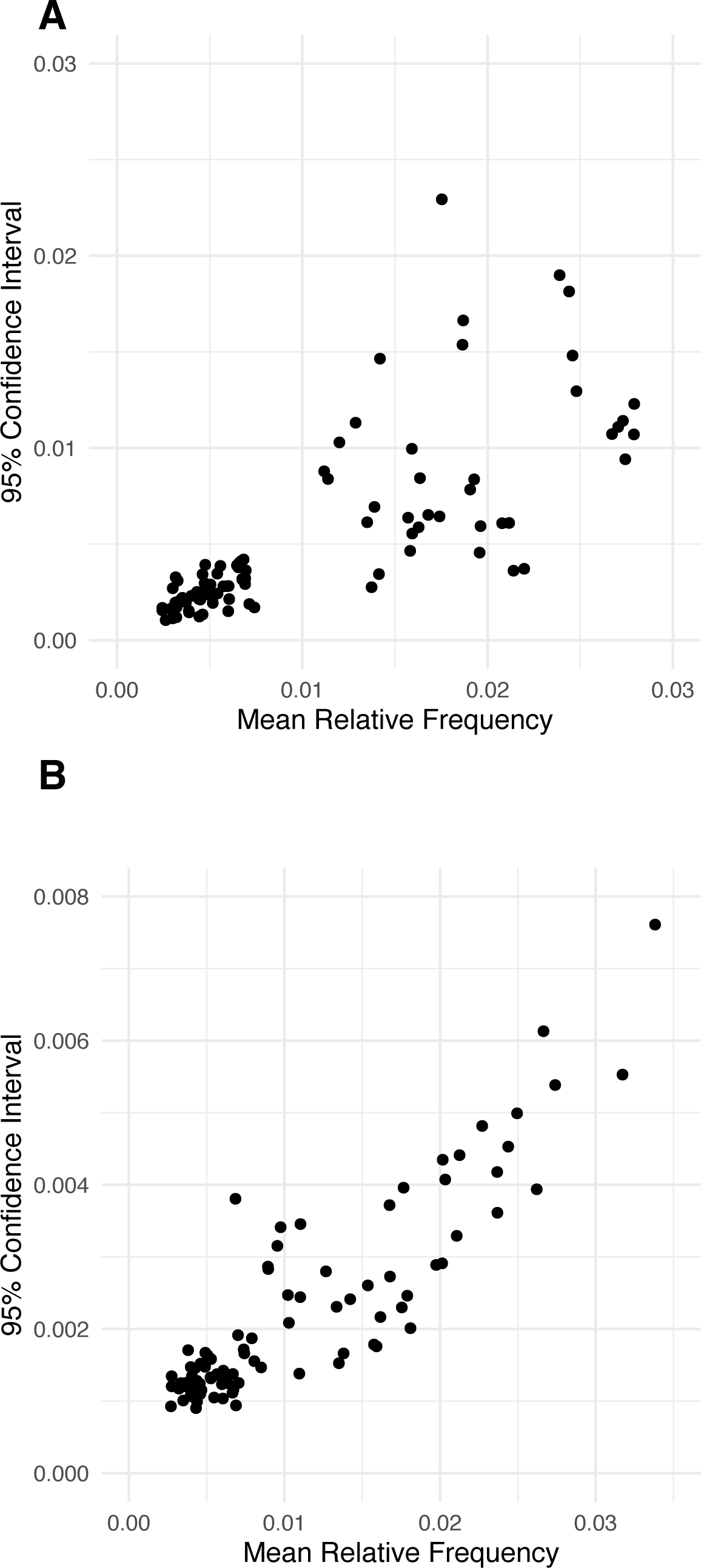
The most frequently mutated motifs tended to show the most variability in mutation frequency when comparing among species. (A) Mean relative frequencies of mutations from the 96-bar graph for the consensus pattern among seven model species (see Figure 3B) were plotted with their corresponding 95% confidence intervals. (B) Mean relative frequencies of mutations from the 96-bar graph for the consensus pattern among 41 species with NCBI SNP data (see Figure 3C) were plotted with their corresponding 95% confidence intervals. For both groups, frequency of mutation at a trinucleotide motif was positively correlated with variability (i.e., larger 95% confidence intervals).

**Fig. S5.**
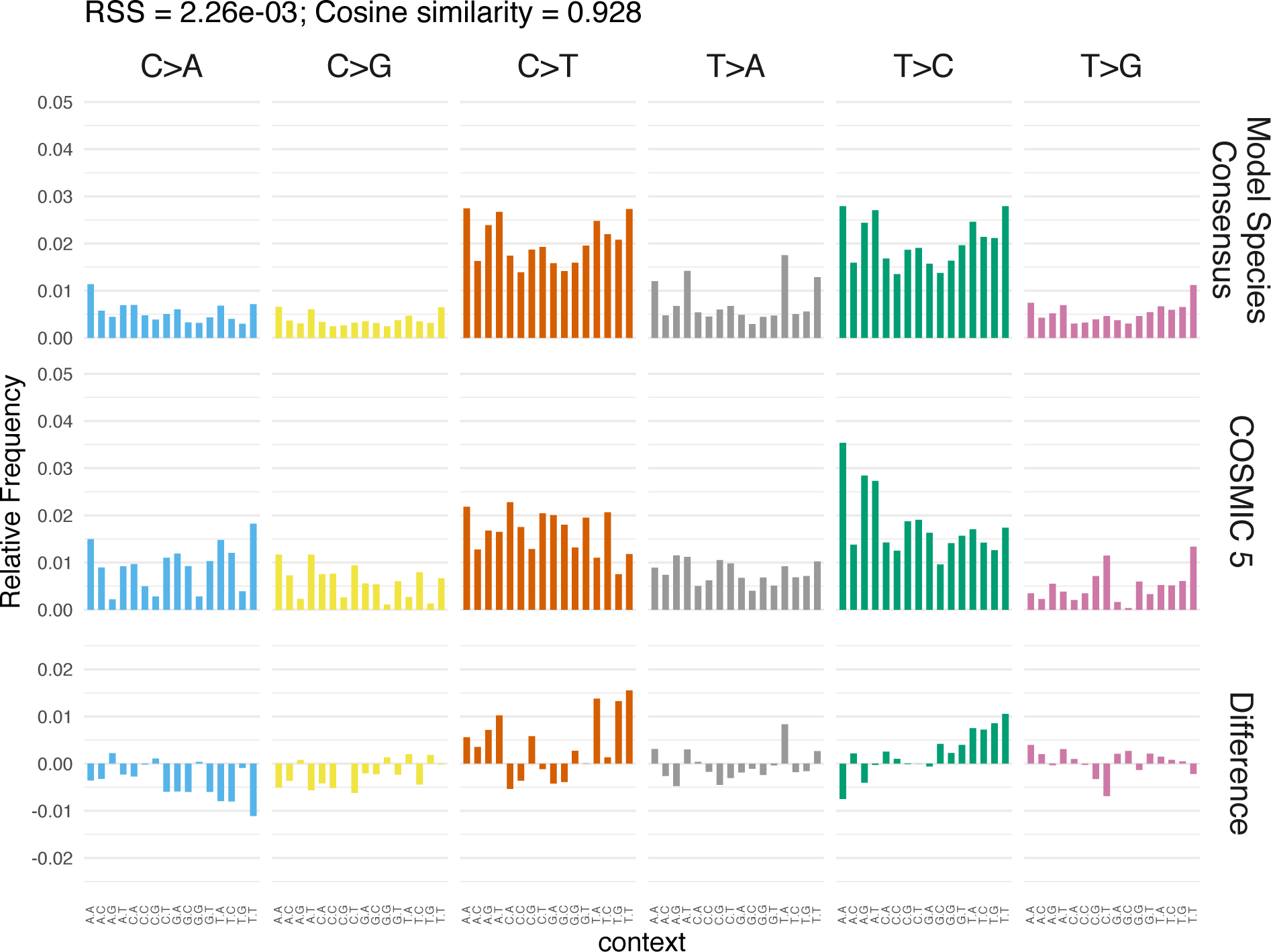
Comparison of the consensus substitution profile mean from seven model species (see Figure 3B) vs. COSMIC signature 5. The cosine similarity value of 0.928 suggested that, despite the evolutionary distance between these model species and human, the underlying biochemical mechanisms creating the intrinsic DNA damage are likely to be well conserved.

**Fig. S6.**
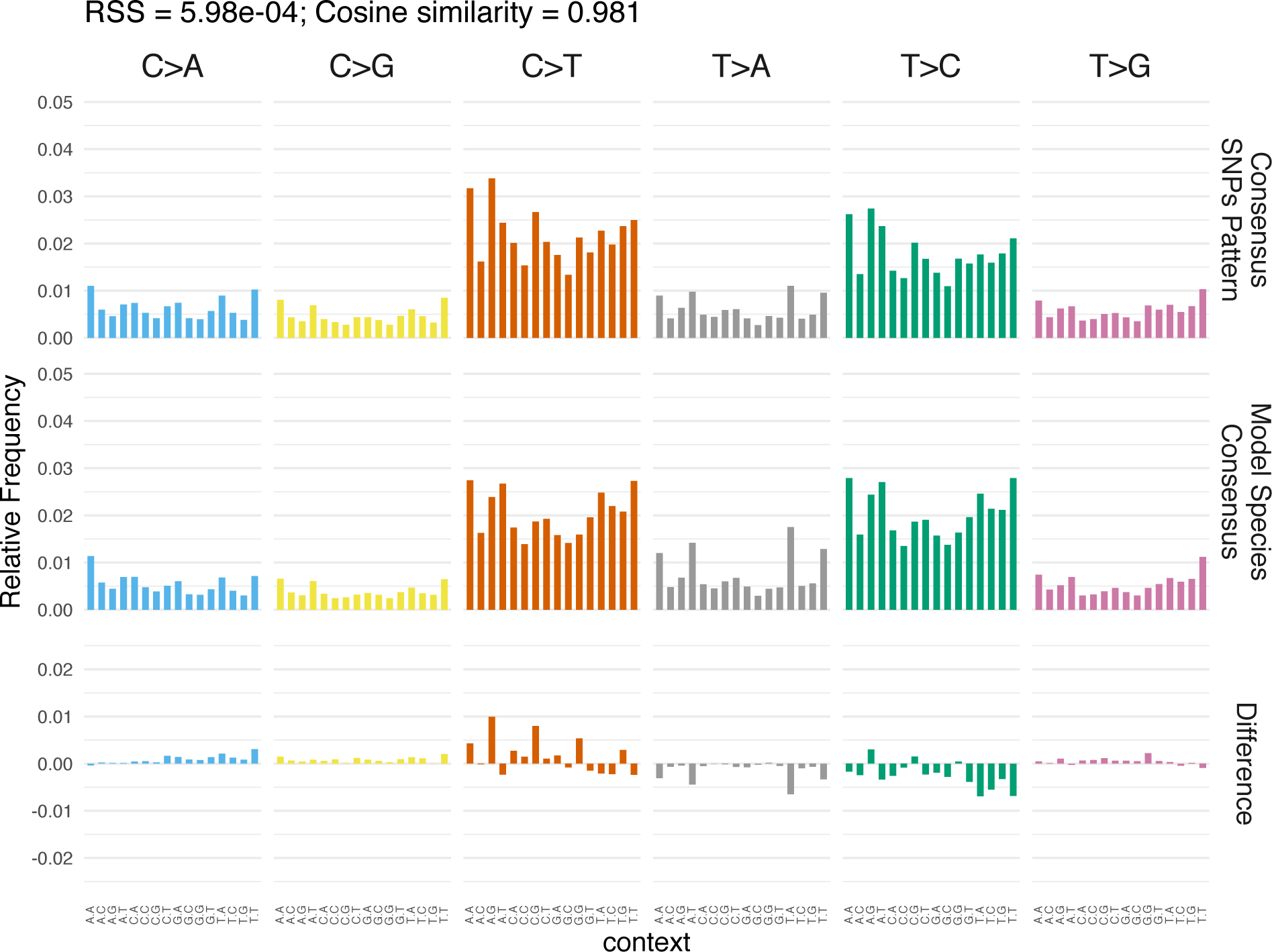
Comparison of the consensus substitution profile mean from 41 (non-human) species with NCBI SNP data (see Figure 3C) vs. the analogous pattern from seven model species (see Figure 3B). The very high cosine similarity value of 0.981 strongly implied that the underlying biochemical mechanisms creating the intrinsic DNA damage are likely to be well conserved across many species.

**Fig. S7.**
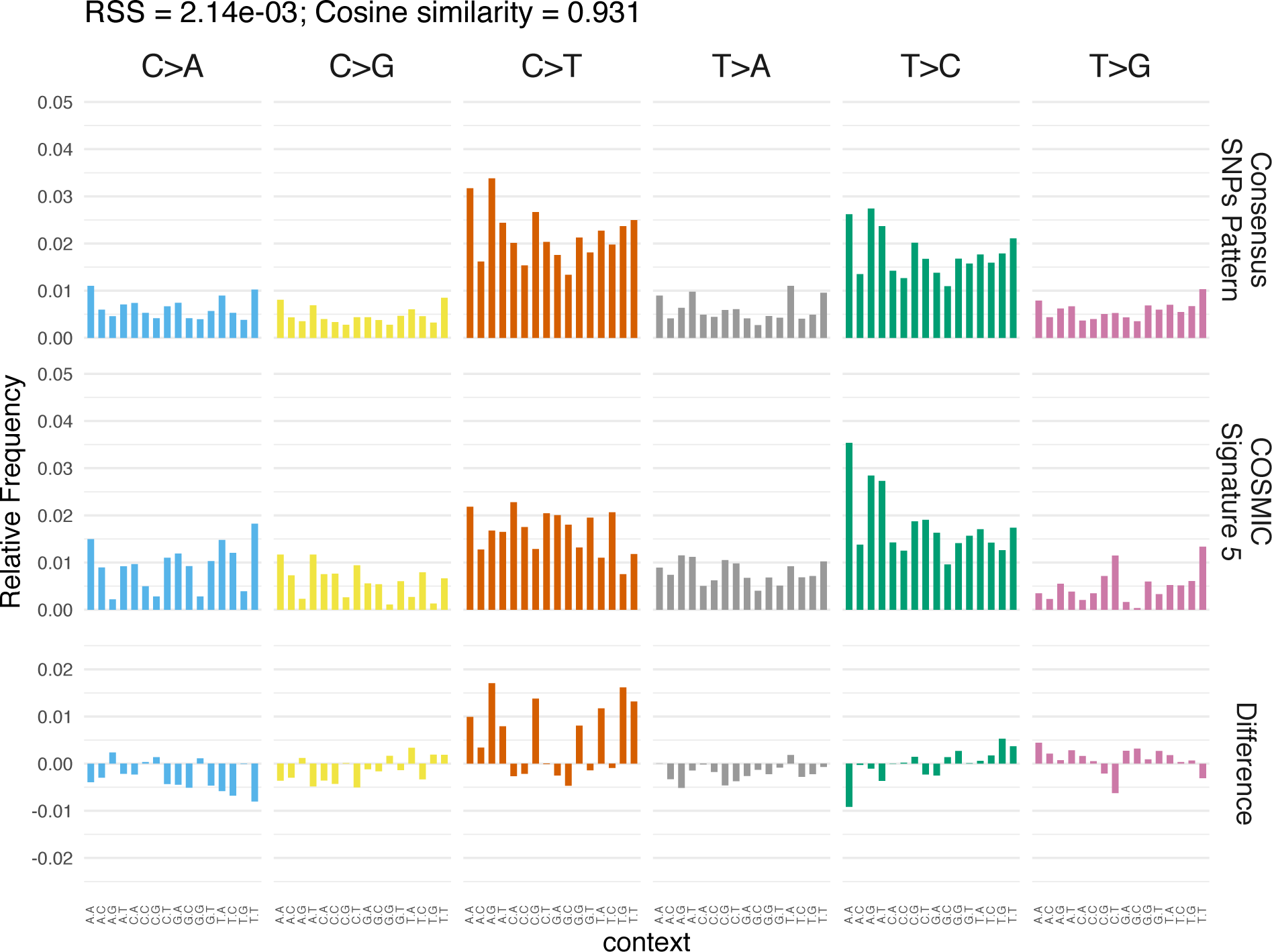
Comparison of the consensus substitution profile mean from 41 (non-human) species with NCBI SNP data (see Figure 3C) vs. COSMIC signature 5. Similar to the model species data, the high cosine similarity value of 0.931 suggested that, despite significant evolutionary divergence vs. human in many species, the underlying biochemical mechanisms which damage the DNA to produce these related substitution patterns are likely to be well conserved.

**Fig. S8.**
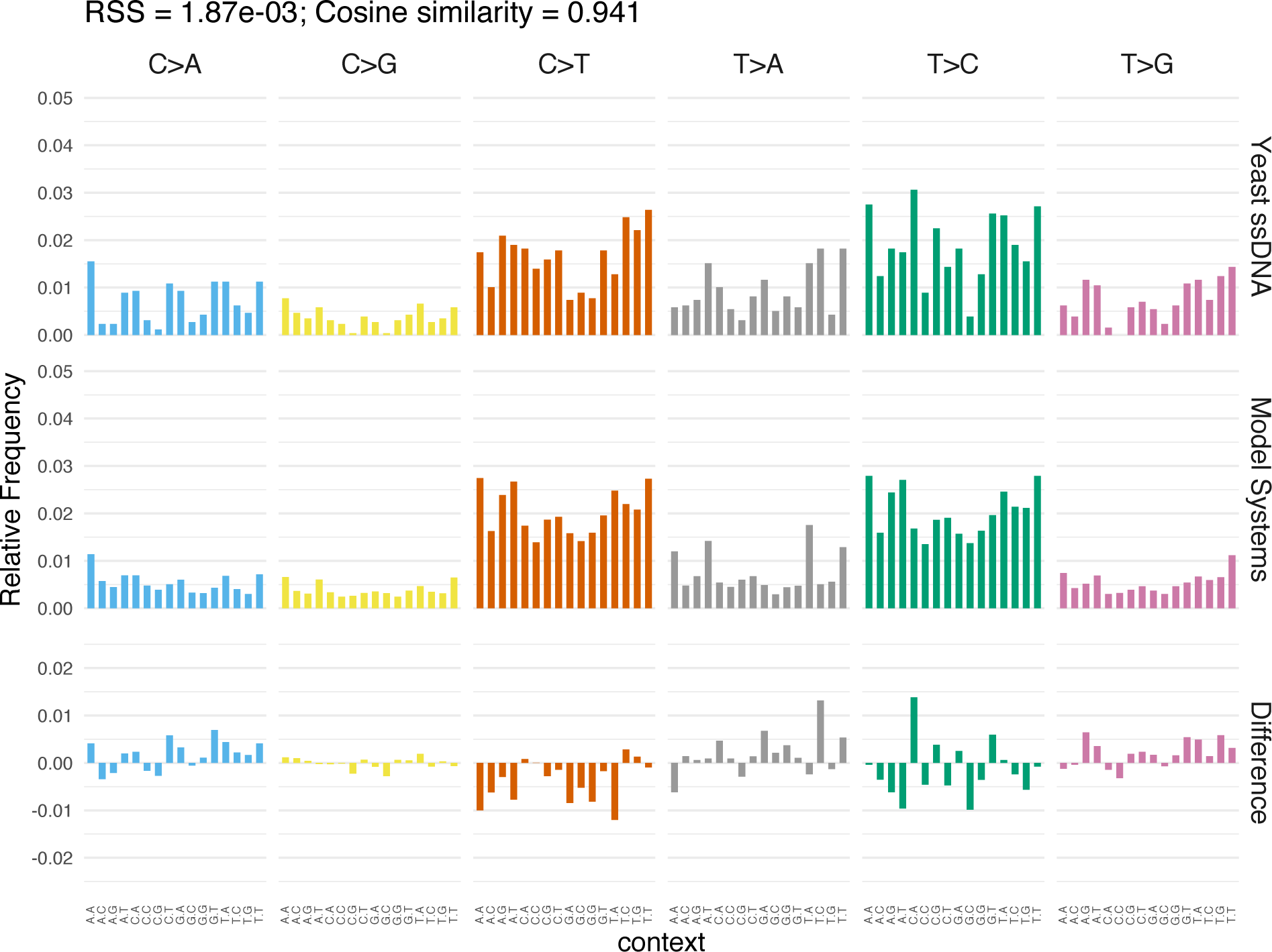
Comparison of the mutational profile observed in yeast single-stranded DNA vs. the consensus pattern among seven model species. Cosine similarity = 0.941.

**Fig. S9.**
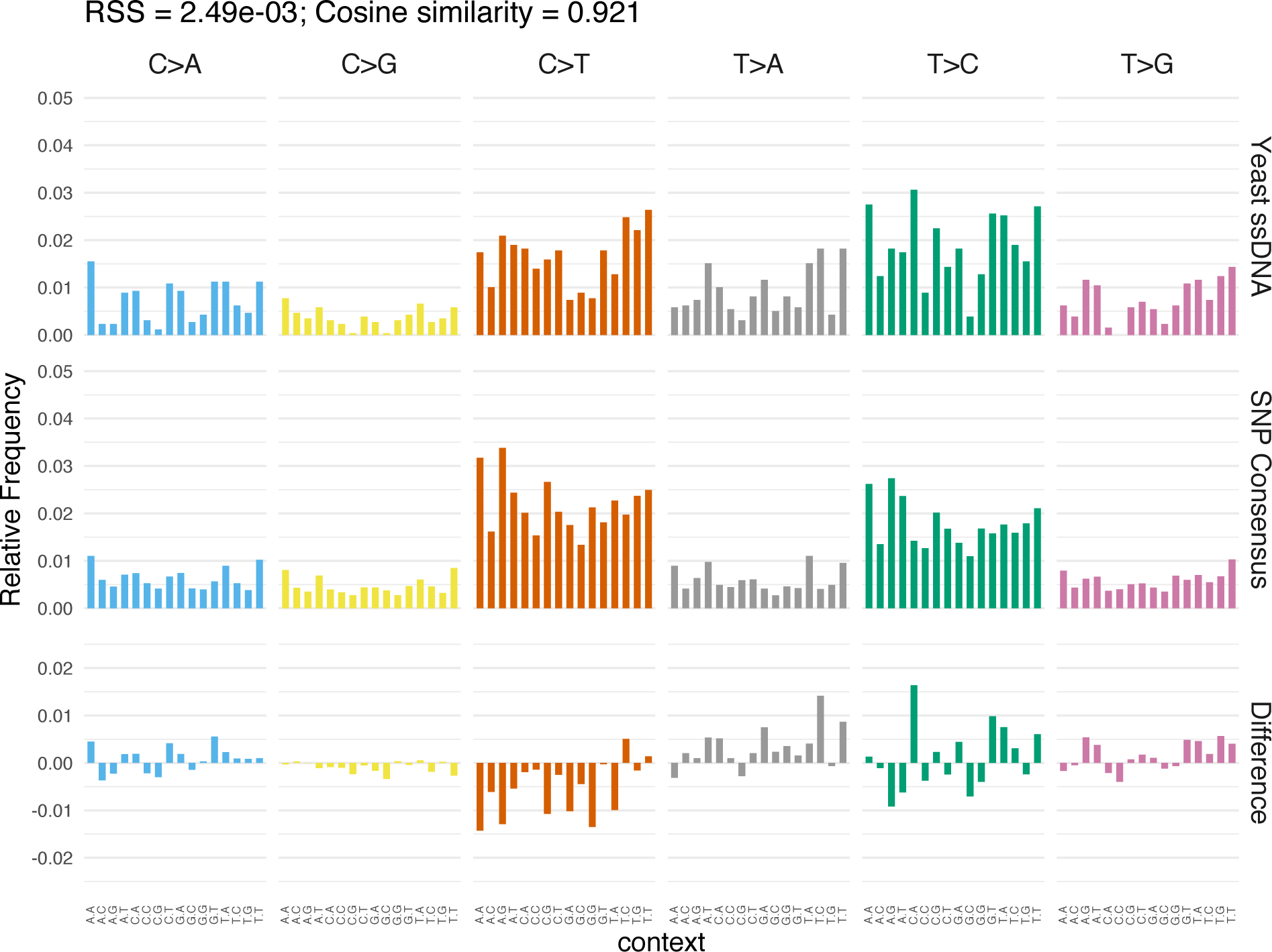
Comparison of the mutational profile observed in yeast single-stranded DNA vs. the consensus pattern from 41 (non-human) species with NCBI SNP data. Cosine similarity = 0.921.

**Fig. S10.**
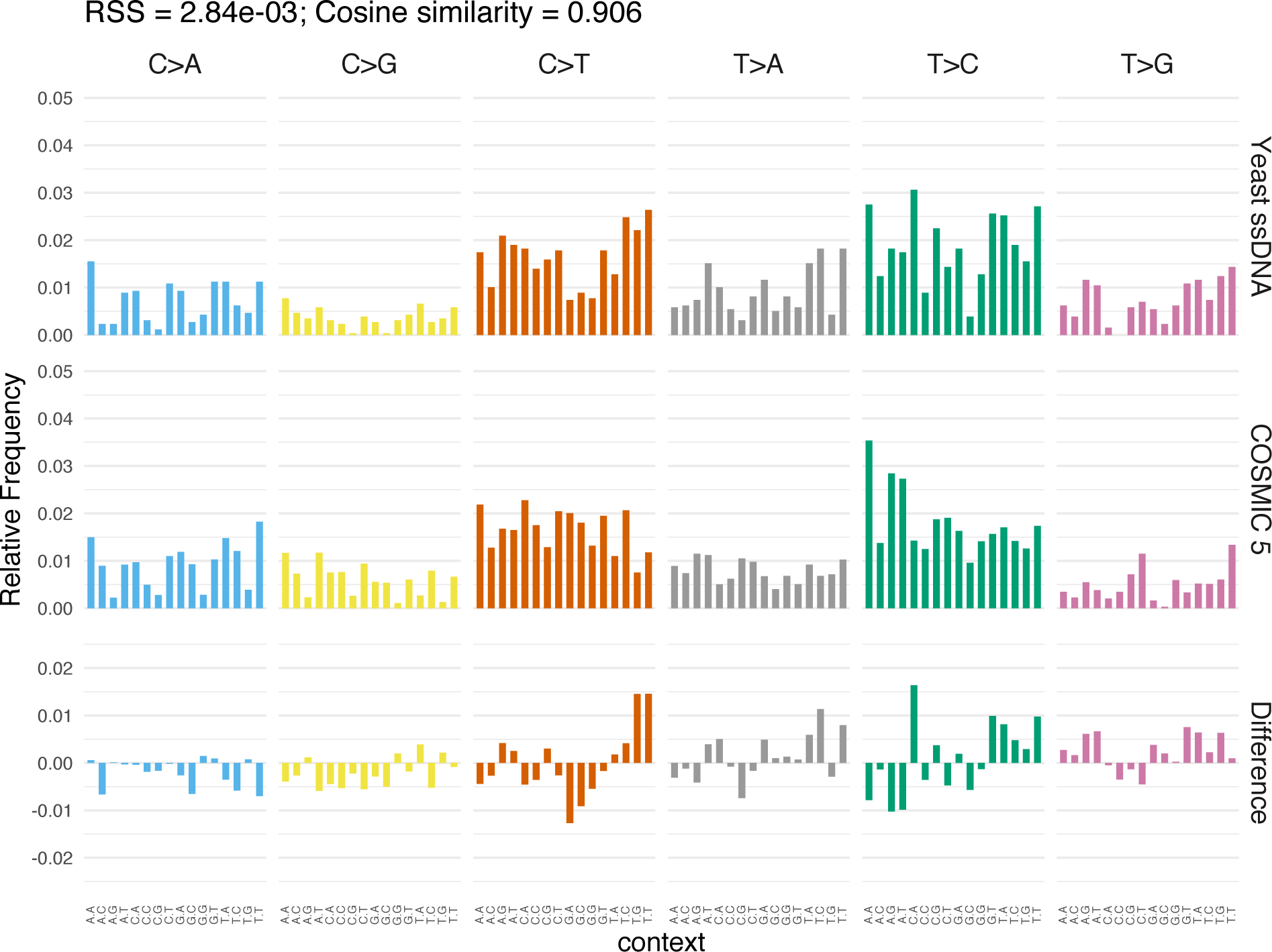
Comparison of the mutational profile observed in yeast single-stranded DNA vs. COSMIC signature 5. Cosine similarity = 0.906.

**Fig. S11.**
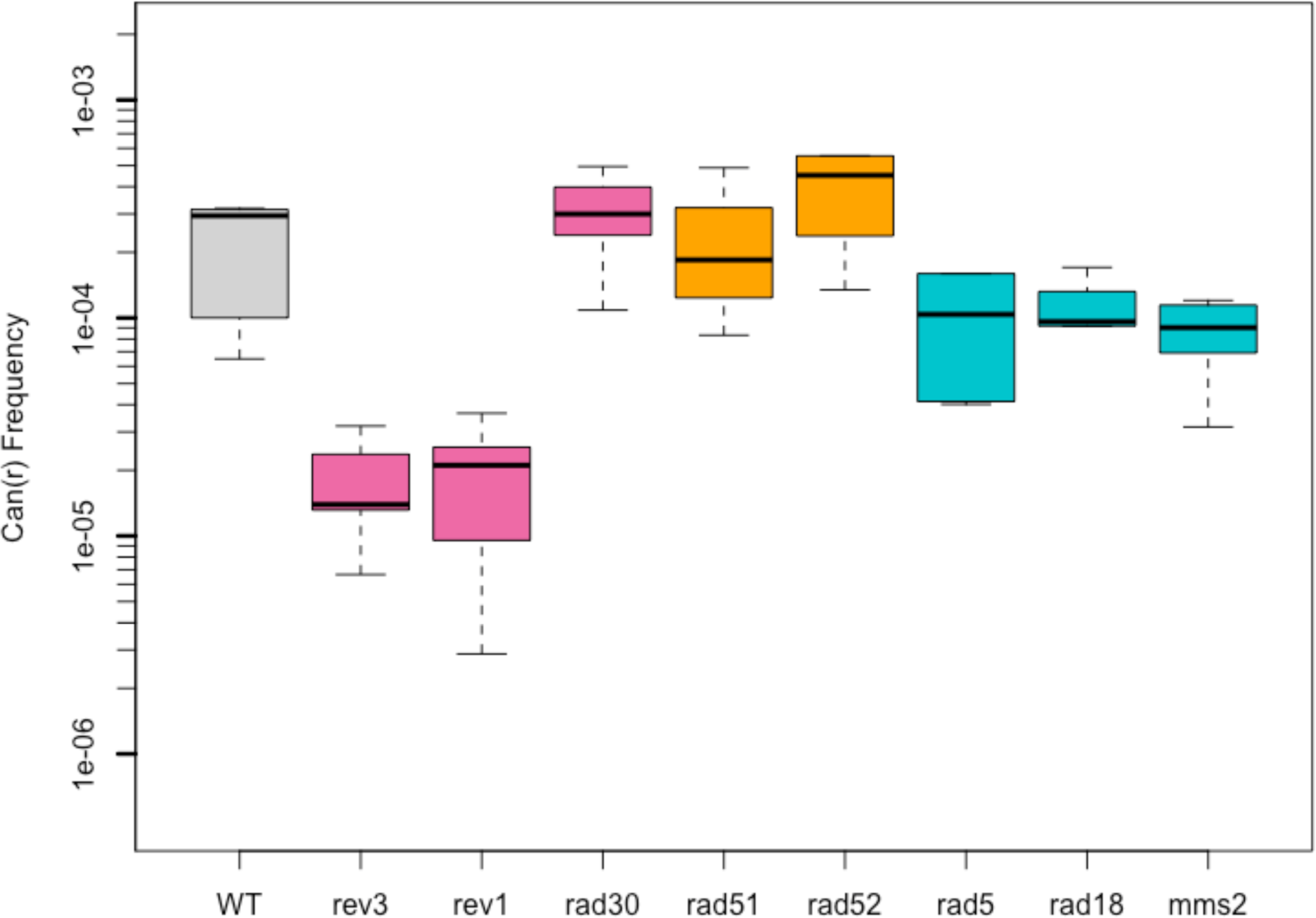
*CAN1* reporter gene inactivation frequency as a function of DNA damage tolerance or DNA repair genotype. Deletion of *REV3* or *REV1* had the largest effect on mutagenesis, as they encode for the catalytic subunit of translesion DNA synthesis (TLS) polymerase zeta and for the sole subunit of a second TLS polymerase, respectively. Deletion of genes involved in error-free damage bypass (*RAD5*, *RAD18*, and *MMS2*) had only minor effects which were not statistically significant, after correction for false discovery for multiple t-tests. Deletion of *RAD30* (TLS polymerase eta) had no effect, nor did deletion of *RAD51* or *RAD52*, both encoding key players in homologous recombination.

**Fig. S12.**
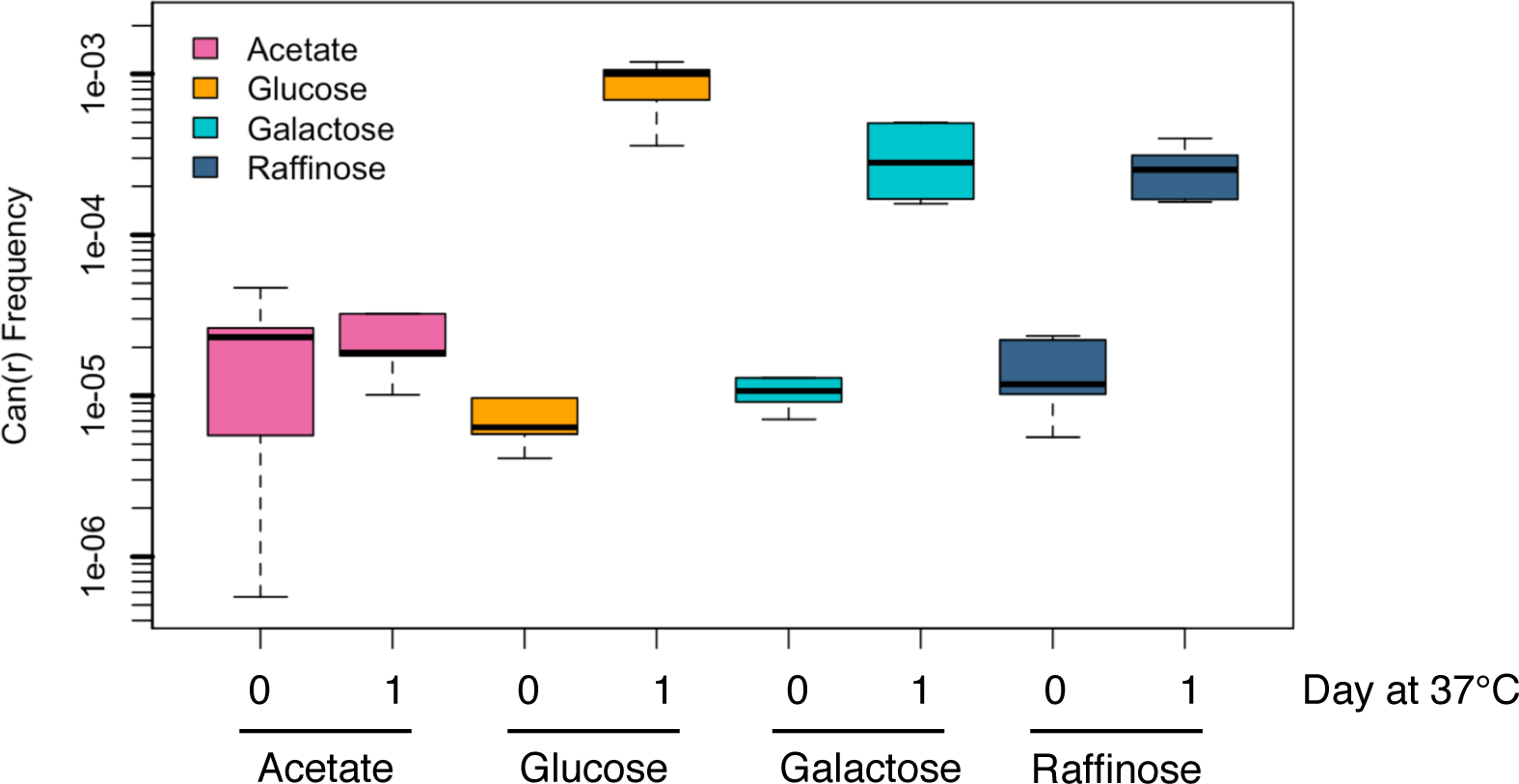
*CAN1* reporter gene inactivation frequency as a function of carbon source. Frequencies at Day 0 denote the background gene inactivation when cells were simply grown at 23°C. Glucose was the most mutagenic, leading to a gene inactivation frequency of ∼10^-3^ after one day at 37°C to damage ssDNA. Acetate, a non-fermentable carbon source that is directly fed into the tricarboxylic acid cycle, then into oxidative phosphorylation, was the least mutagenic. Galactose and raffinose, which have mixed “respiro-fermentative” metabolism, were intermediate in mutagenicity.

**Fig. S13.**
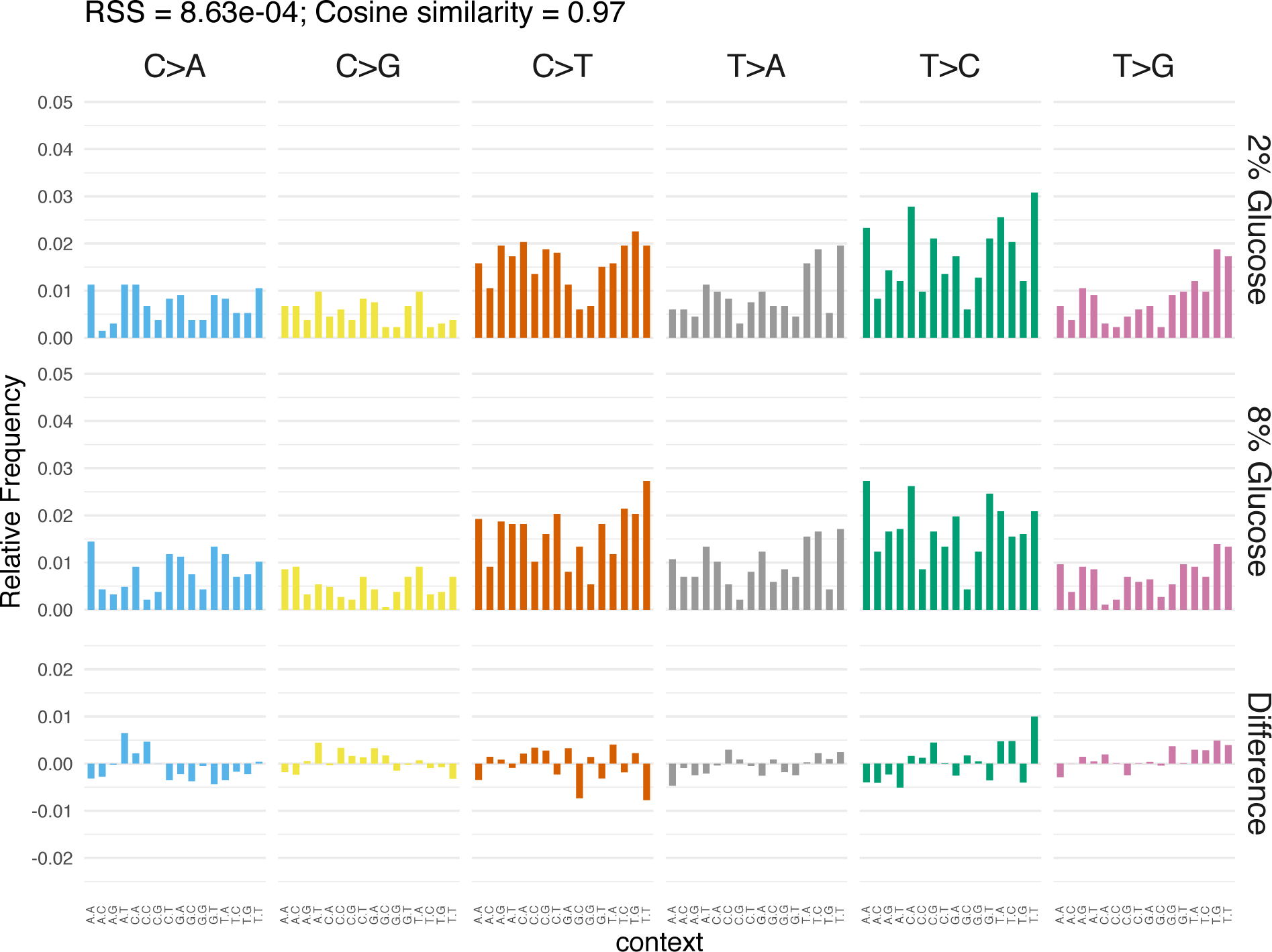
Comparison of the patterns from metabolizing 2% glucose vs. 8% glucose revealed a very high cosine similarity (0.970), indicating that in 8% glucose, cells simply acquired more of the same pattern of mutations.

**Fig. S14.**
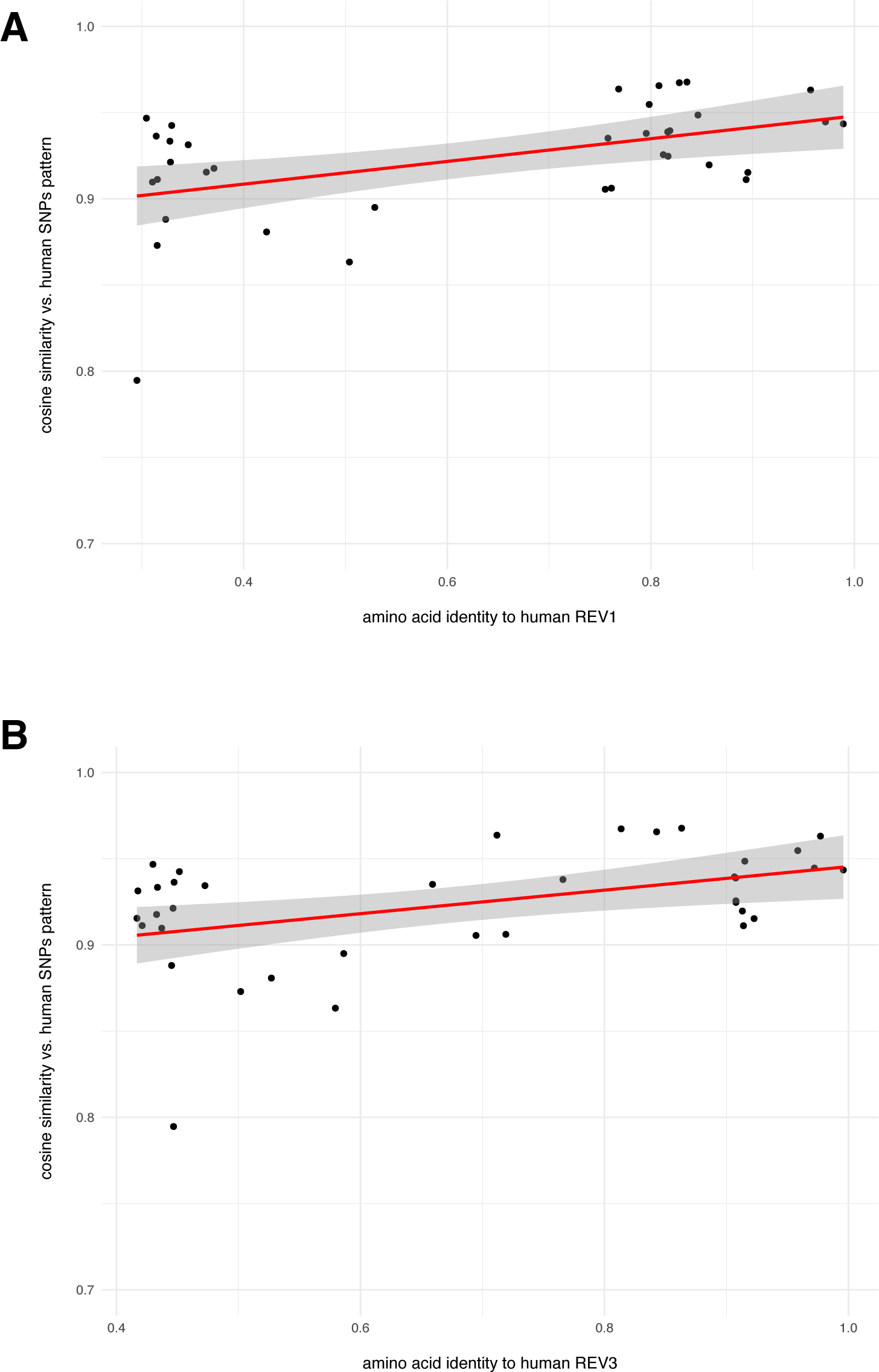
Cosine similarity, when comparing base substitution patterns in (non-human) NCBI SNP species vs. NCBI human SNPs pattern, was plotted as a function of amino acid identity to the two key translesion DNA synthesis polymerases REV1 (A) and REV3 (B). 95% confidence intervals are shaded in gray. Species whose REV1 and REV3 homologs are more similar to the human polymerases tended to have base substitution patterns with higher cosine similarity. Remarkably, many species with relatively low amino acid identity (less than 0.5) still had cosine similarity values > 0.9, underscoring the extensive conservation of COSMIC 5-like patterns throughout evolution.

**Fig. S15.**
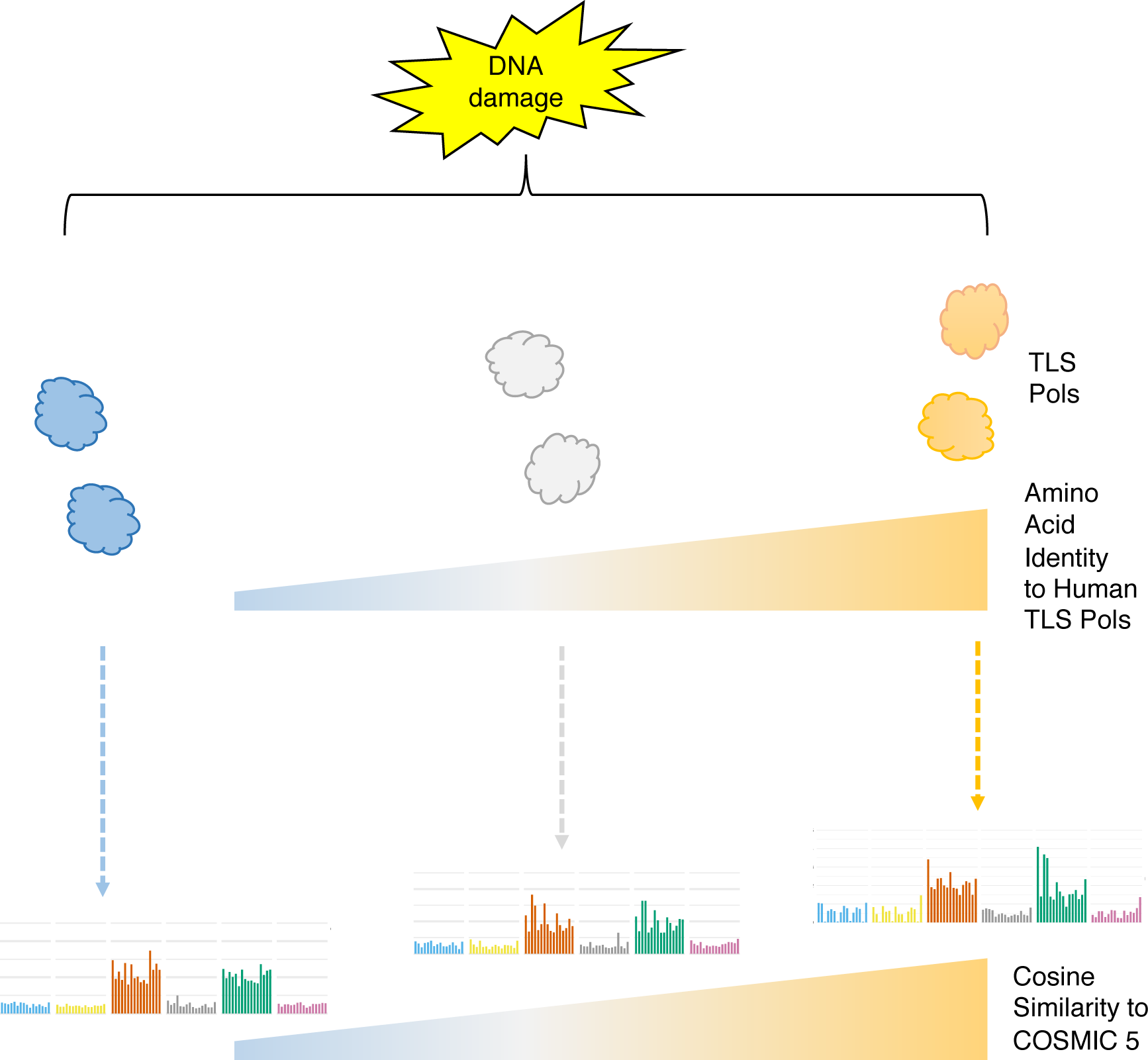
Model explaining how a continuum of COSMIC 5-like patterns arises among diverse species. DNA damage from metabolic source(s) are similar among species. Translesion DNA synthesis polymerases (TLS Pols) insert bases in an error-prone manner opposite the base damage. The species with TLS polymerases which are most similar to human (shown in orange) produce intrinsic base substitution patterns with the highest cosine similarity values (∼ 0.97) when compared to COSMIC 5. Species with TLS polymerases that are most diverged from those in human (shown in blue) yield base substitution patterns in the lower end of the cosine similarity range (∼ 0.90) vs. COSMIC 5. Species with TLS polymerases that are intermediate also tend to produce base substitution patterns that are intermediate in cosine similarity. Notably, even the most diverged TLS polymerases can still yield cosine similarity values > 0.9 vs. COSMIC 5.

